# Leveraging WNT Hyperactivation to Kill Colorectal Cancer While Rejuvenating Healthy Intestine

**DOI:** 10.1101/2025.10.05.680591

**Authors:** George Eng, Jonathan Braverman, Manish Gala, Omer Yilmaz

## Abstract

WNT signaling maintains intestinal homeostasis yet drives colorectal cancer (CRC) when constitutively activated by APC mutations. We overturn the fundamental assumption that APC-mutant tumors exist at maximal WNT activation, revealing instead that cancer cells occupy a precarious "WNT-just-right" zone along a signaling continuum. This discovery exposes an unprecedented therapeutic vulnerability: while normal intestinal epithelium thrives with enhanced WNT signaling, APC-mutant tumor cells undergo apoptosis when pushed beyond their oncogenic setpoint, a phenomenon we term "over-WNTing." Through systematic organoid-based screening, we identified that WNT hyperactivation through multiple approaches: GSK3 inhibition, concentrated WNT proteins, or APC knockdown in GSK3-null backgrounds, selectively kills tumor cells by hyperactivating the driving pathway of CRC. Mechanistically, over-WNTing in APC-mutant cells triggers spillover into non-canonical planar cell polarity signaling, where RHOC upregulation induces ROCK1/2-mediated apoptosis. We demonstrate therapeutic efficacy across the neoplastic continuum, from adenomas to metastatic CRC, including patient-derived tumors, validating GSK3 inhibition with a novel nanoparticle formulation. This discovery enables the first cancer therapy that simultaneously enhances normal tissue function while eliminating tumors. "Over-WNTing" effectively treats adenomas and both mouse and patient-derived CRC, establishing a therapeutic paradigm that exploits fundamental differences in cellular WNT biology to achieve the dual benefit of eliminating cancer while promoting healthy tissue regeneration.

## INTRODUCTION

WNT signaling orchestrates intestinal homeostasis through precise regulation of stem cell self-renewal and differentiation.^1–4^. When constitutively activated—most commonly through APC mutations— this same pathway drives adenoma formation and progression to colorectal cancer (CRC).^5^ This duality has created a vexing therapeutic paradox: the pathway essential for normal tissue maintenance also drives and maintains malignancy in over 90% of colorectal cancers. For decades, this paradox has rendered WNT signaling clinically intractable, with every therapeutic strategy focused on pathway inhibition failing due to devastating on-target toxicity in healthy stem cells, that depend on WNT signaling for survival and function.^6–13^ These failures arose from a core assumption that APC-mutant tumors, having lost a key negative regulator, exist at maximal WNT activation and therefore would be treatable by WNT suppression.

Here, we investigate this paradox by overturning this fundamental assumption. Rather than viewing APC loss as inducing maximal WNT activation that must be reduced, we demonstrate that cancer cells exist in a precarious WNT-optimal state—a “just-right” signaling window between their baseline constitutive activation and a lethal threshold.^14–19^ This positioning creates an unexpected vulnerability: while normal intestinal cells operating at physiological WNT levels tolerate substantial pathway enhancement, APC-mutant cancer cells are inhibited with any further WNT activation.

Through systematic organoid-based screening validated across patient-derived tumors and in vivo models, we used multiple orthogonal approaches to exploit this vulnerability through three strategies: pharmacologic intervention (selective GSK3 inhibition with LY2090314), protein-based activation (concentrated WNT3a/R-spondin), and genetic manipulation (disruption of negative regulators). Using quantitative WNT reporters, we provide direct molecular evidence that APC-mutant cancer cells maintain sub-maximal pathway activation and can be pushed beyond their viable range—a phenomenon we term "over-WNTing." This hyperactivation state revealed an unexpected mechanistic basis for cancer-selective toxicity.

Transcriptomic analysis of “over-WNTed” of colon cancer organoids uncovered aberrant activation of non-canonical planar cell polarity signaling as the causal molecular mechanism. Specifically, we identified RHOC upregulation as the critical mediator, with beta-catenin directly inducing RHOC transcription at supraphysiological WNT levels, leading to ROCK1/2-mediated apoptosis. Genetic deletion of RHOC or ROCK1/2, completely prevented over-WNTing-induced death, establishing this axis as necessary for the selective toxicity.

We validate this therapeutic principle across multiple disease models from adenomas to metastatic cancer using novel nanoparticle formulations that achieve 93% tumor reduction—exceeding standard chemotherapy—while exhibiting no toxicity and enhancing intestinal epithelial regeneration. This work fundamentally reframes oncogenic signaling from intractable driver pathways to exploitable vulnerabilities. By demonstrating that constitutive oncogenic activation creates inflexible cell states susceptible to further pathway activation, we establish the first therapeutic paradigm where cancer treatment rejuvenates rather than compromises normal tissue function.

## RESULTS

### GSK3 inhibition is a potent WNT agonist in the intestine

Canonical WNT signaling is initiated by extracellular WNTs and RSPO1/2/3/4, which bind to Frizzled and Lgr4/5 receptors to inhibit the beta-catenin destruction complex.^1,20,21^ Subsequently, beta-catenin translocates to the nucleus, binds to LEF/TCF, and induces transcriptional changes.^22,23^ Since GSK3 alpha and GSK3 beta along with APC, participate in the beta-catenin destruction complex, we hypothesized that pharmacologic inhibition of GSK3 could phenocopy *APC* mutations, inducing growth in the absence of exogenous WNT. ^12,24,25^ (Schematic 1) The accompanying WNT signaling diagram is an abbreviated depiction of the specific WNT pathway elements modulated in this study, and many other WNT pathway elements are also important but not highlighted.

We tested commercially available small molecule inhibitors of GSK3 alpha and beta for their capacity to support the growth of primary wildtype mouse colonic epithelium as organoids in the absence of other exogenous WNT ligands (Fig 1A). GSK3 alpha and beta have a high degree of homology and GSK3 inhibitors generally act on both isozymes.^26–28^ Only LY2090314 permitted organoid growth over a wide concentration range and demonstrated dose-dependent capacity to grow primary colon organoids (Fig 1A,B).^29^ Quantitative TOP/Flash reporter assays confirmed robust, dose-dependent WNT pathway activation, with LY2090314 demonstrating higher absolute induction than standard WRN recombinant WNT factor supernatant media across nearly a thousand-fold dilution range (Fig. S1A,B).^30,31,32^ While conventional drug discovery approaches that measure IC50 values against purified GSK3 proteins would predict CHIR99021 and LY2090314 to be similarly potent, our intact organoid system revealed that only LY2090314 was capable of functional activity in the intestinal organoids.^33–35^ This finding highlights the power of organoid-based functional screening to capture physiologically relevant drug activities.

**Figure 1.**
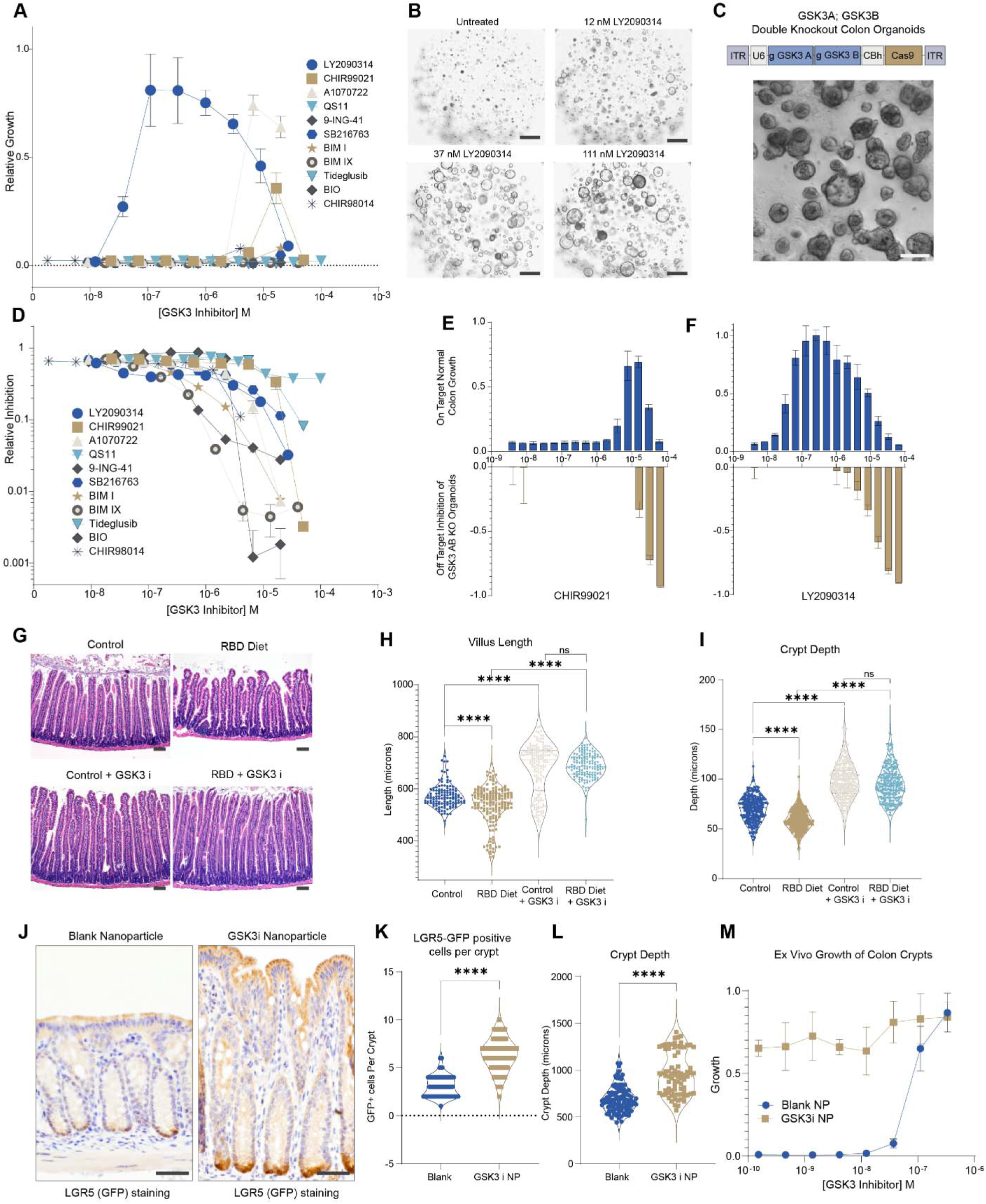
GSK3 Inhibition as a Potent WNT Agonist for Intestinal Regeneration. (A) GSK3 inhibitor screen for wildtype mouse colon organoid growth measured by resazurin metabolic activity assay after 4 days in culture. Eleven small molecule GSK3 inhibitors were tested for their capacity to support wildtype mouse colon organoid growth in media lacking WNT3A or R-spondin. Growth is expressed as fold-change relative to untreated control organoids. The decrease at higher concentrations reflects compound toxicity at supraphysiological doses. n=3 biological replicates per condition. (B) Representative brightfield images of wildtype mouse colon organoids grown with LY2090314 (1 μM) without additional WNT agonists, showing cystic morphology characteristic of WNT-activated organoids similar to APC-mutant organoids. Images taken after 4 days in culture. (C) Schematic of CRISPR-Cas9 dual-guide vector construct used to generate GSK3A/B double knockout organoids, and representative brightfield image demonstrating their growth in WNT agonist-free media, confirming genetic WNT independence. (D) GSK3 inhibitor screen for off-target growth effects using GSK3A/B knockout organoids with the same compounds tested in (A). Growth measured by resazurin assay demonstrates compound specificity. n=3 biological replicates per condition. (E,F) On-target versus off-target validation comparing effects of CHIR99021 (E) and LY2090314 (F) on wildtype organoids (top graphs, demonstrating WNT activation) versus GSK3A/B knockout organoids (bottom graphs, revealing off-target effects). LY2090314 shows superior on-target selectivity with minimal off-target toxicity. (G) Hematoxylin and eosin histology of small intestine from mice fed control diet or regional basic diet (RBD) to induce environmental enteropathy, with or without GSK3 inhibitor nanoparticle treatment (daily enemas for 2 weeks). GSK3 inhibitor treatment restores villus architecture and crypt depth in enteropathy model. n=5 mice per group, representative of two independent experiments. (H,I) Quantification of villus length (H) and crypt depth (I) from histology in (G), demonstrating GSK3 inhibitor-mediated intestinal regeneration in the enteropathy model. (J) Immunohistochemistry for GFP in LGR5-EGFP-CreERT2 mice after 2 weeks of enema treatment with blank nanoparticles or GSK3 inhibitor nanoparticles (3 times weekly). GSK3 inhibitor treatment increases LGR5+ stem cell numbers and crypt depth. n=5 mice per group. (K,L) Quantification of LGR5-GFP positive cells per crypt (K) and crypt depth (L) from immunohistochemistry in (J), confirming enhanced stem cell function and tissue regeneration. (M) Ex vivo organoid growth potential of isolated colon crypts from mice in (J), cultured in dose-response of GSK3 inhibitor. Crypts from GSK3 inhibitor-treated mice show enhanced growth capacity, demonstrating improved stem cell function. Crypts isolated 24 hours after final treatment. Growth measured by resazurin assay after 4 days. Statistical Analysis: Data presented as mean ± SD. Statistical comparisons performed using unpaired two-tailed t-tests for (H, I, K, L) and one-way ANOVA with Tukey’s post-hoc test for dose-response curves (A, D, M). ****p<0.0001, ***p<0.001, **p<0.01, *p<0.05. Scale Bars: 500 μm (B), 250 μm (C), 100 μm (G, J). GSK3i refers to LY2090314 throughout. See also Figure S1.

**Figure S1:**
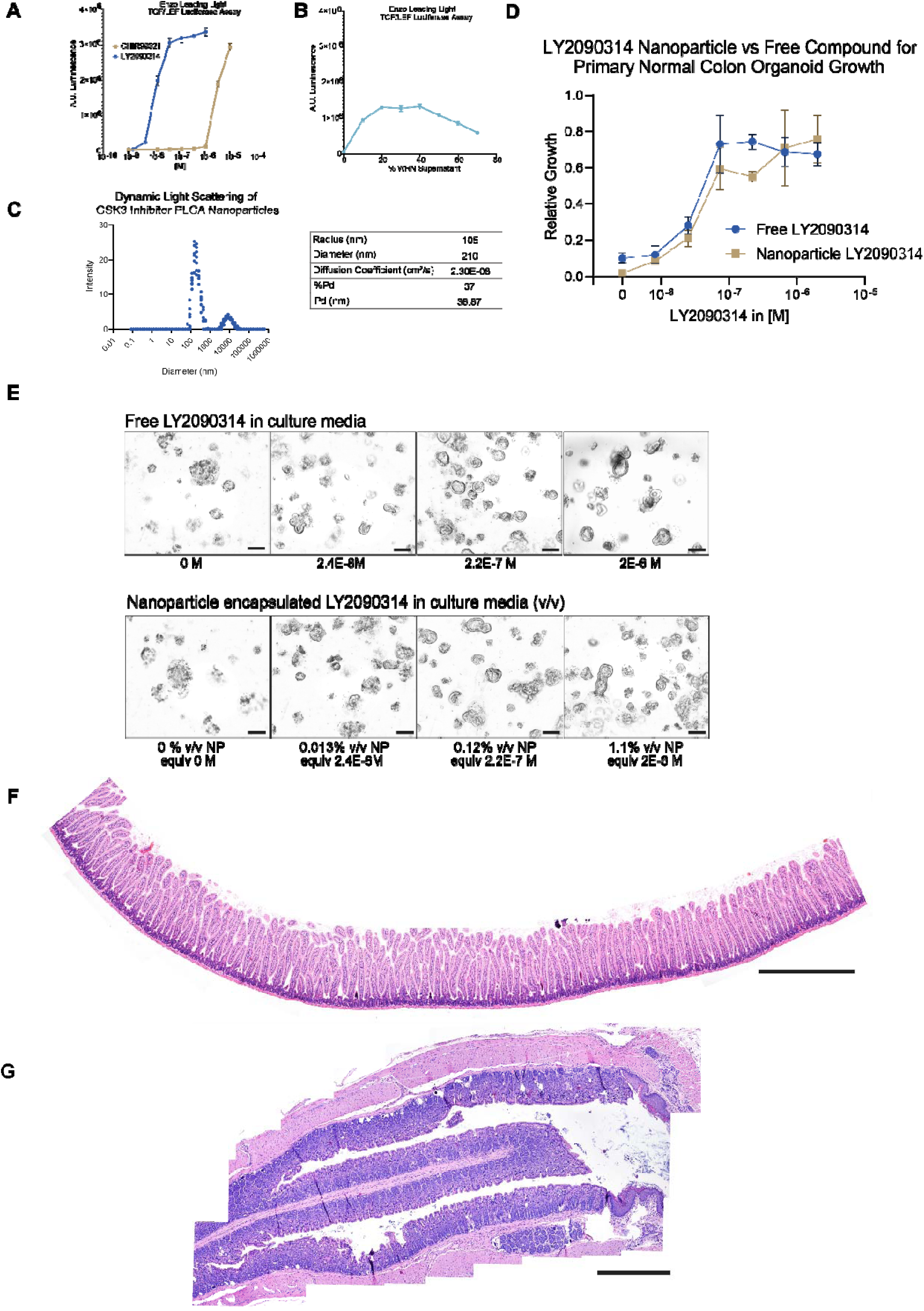
Characterization of LY2090314 WNT activation, and validation of PLGA nanoparticle encapsulation of LY2090314. (A,B) Enzo Leading Light TCF/LEF luciferase reporter assay measuring WNT pathway activation. (A) Dose-response curves comparing LY2090314 and CHIR99021. (B) Dose-response curve of WNT3A-conditioned media (WRN) as a positive control. Data shown as mean ± SD, n=3 biological replicates. (C) Dynamic light scattering analysis of PLGA nanoparticles containing LY2090314. Size distribution shows mean diameter of 210 nm with polydispersity index (PDI) of 38.87%. (D) Comparison of wildtype mouse colon organoid growth with free LY2090314 versus PLGA nanoparticle-encapsulated LY2090314. Growth was quantified by resazurin metabolic assay measuring viable cell mass after 72 hours of culture. Data normalized to the highest growth condition, shown as mean ± SD, n=4 biological replicates. (E) Representative brightfield microscopy images of organoids from (D) after 72 hours of culture at indicated concentrations. Scale bar: 200 μm. (F) Representative stitched image of small intestine from RBD mouse following 7 days of daily oral gavage with LY2090314 nanoparticles (13 consecutive 20x fields). Analysis of 2,196 crypts across 5 treated mice revealed enhanced villus height and crypt depth consistent with WNT-mediated regeneration, with zero adenoma formation. The physiological hyperplasia observed reflects enhanced epithelial growth without neoplastic transformation. Scale bar = 1 mm. (G) Representative stitched image of colon from mouse following 2 weeks of enema delivery with LY2090314 nanoparticles (3 treatments per week, 25 consecutive 20x fields). Comprehensive analysis of 1,664 colonic crypts across 3 treated mice revealed enhanced crypt depth and epithelial thickness as expected from WNT activation, with zero adenoma formation or aberrant crypt foci. The regenerative hyperplasia observed is consistent with enhanced stem cell function without neoplastic transformation. Scale bar = 1 mm.

Recent structural biology work provides mechanistic insight into why the living tissue context matters so profoundly for GSK3 function.^36^ Even when phosphomimetic mutations perfectly recapitulate the structural positioning of phosphorylated substrates in crystal structures, they exhibit 1000-fold reduced activity compared to native phosphopriming—revealing that catalytic efficiency requires dynamic molecular interactions beyond just GSK3 and beta-catenin. In living tissue models such as organoids, GSK3 functions within the Axin/APC scaffolding complex and CK1-mediated phosphopriming of beta-catenin at Ser45 enables processive phosphorylation. This tissue-specific regulatory dependency explains why our organoid-based functional screen successfully identified LY2090314 as uniquely capable of sustaining WNT pathway activation: it effectively engages GSK3 within its native phosphopriming and scaffolding environment. The approach demonstrates how intact tissue systems can reveal functional drug properties that biochemical assays measuring isolated enzyme activity cannot predict, providing a more accurate and physiologically relevant path for WNT pathway modulator discovery.

**Schematic 1:**
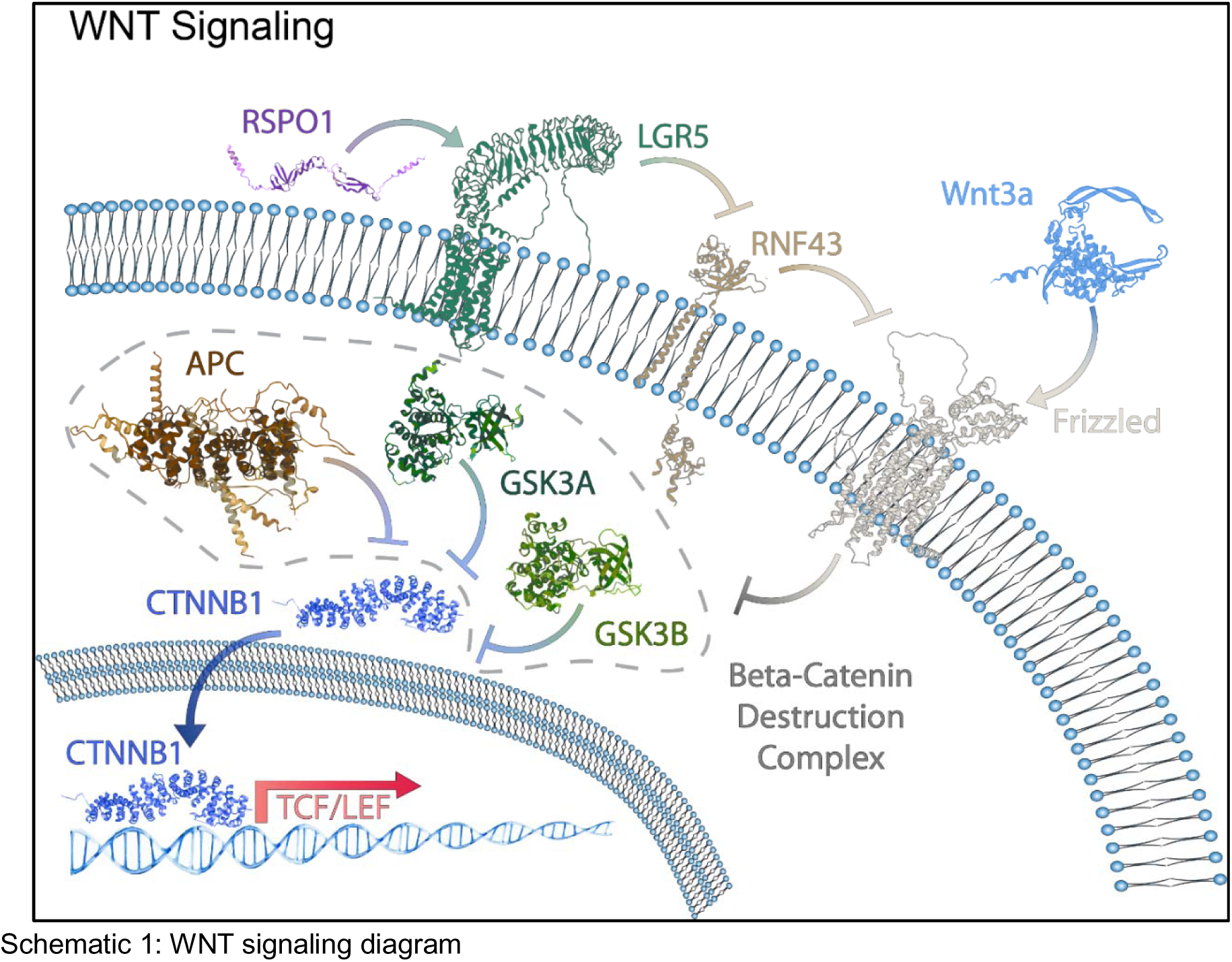
**WNT signaling diagram**

To rigorously evaluate GSK3 inhibitor specificity, we engineered the first colon organoids genetically deficient in both GSK3α and GSK3β using CRISPR-Cas9 technology. These double knockout organoids exhibit constitutive WNT activation and grow independently of exogenous WNT ligands, phenocopying APC loss-of-function mutations for niche independence (Fig 1C)^18^. By eliminating the target proteins, this system definitively distinguishes on-target from off-target effects—any growth inhibition in these cells must arise from off-target activities since both isoforms of GSK3 are absent. Testing the same panel of GSK3 inhibitors on these double knockout organoids revealed a striking pattern (Fig 1D). Our integrated analysis—plotting on-target growth promotion against off-target toxicity— revealed that most GSK3 inhibitors are functionally non-selective in intestinal tissue (Fig 1D-F). LY2090314 emerged as uniquely selective, maintaining activity across a thousand-fold concentration range (Fig 1F), while CHIR99021 and other inhibitors showed substantial off-target toxicity at biologically active concentrations (Fig 1E)^37–40^ This discovery reconciles why GSK3 inhibitors have failed as WNT activators: the limitation arises from inadequate compound selectivity, not target biology. The ability of a single selective inhibitor to completely replace exogenous WNT proteins validates both GSK3’s centrality to WNT signaling and the critical importance of tissue-specific screening in drug discovery.

We validated the *in vivo* effects of GSK3 inhibition by developing nanoparticle-encapsulated LY2090314 using poly-lactic-co-glycolic acid polymeric nanoparticles which protect the compound from degradation and early absorption in the digestive tract (Fig. S1C).^35,41^ Quantitative dose-response analysis demonstrated that nanoparticle-encapsulated LY2090314 maintained identical growth-promoting activity to free drug in normal mouse colon organoid cultures, with corresponding morphological equivalence, confirming both preserved bioactivity following encapsulation, and no specific activity added by encapsulation (Fig. S1D,E).

We, then, queried in vivo efficacy using a mouse model of environmental enteropathy, a condition affecting millions globally where nutritional insufficiency induces intestinal epithelial damage and crypt atrophy. ^42^ This disease model demonstrates that GSK3 inhibition—which previously was only used as a research tool—could function as a direct, pharmacologically tractable method to enhance normal intestinal growth. Mice fed a regionally basic diet recapitulate the human disease phenotype, exhibiting characteristic villus shortening and crypt atrophy (Fig 1G). ^43–46^ The nanoparticle-delivered GSK3 inhibitor demonstrated safe, effective pharmacological enhancement of normal intestinal tissue function. Treatment of healthy control mice demonstrated that GSK3 inhibition could augment normal intestinal growth establishing the first clear evidence that selective GSK3 inhibition functions as a direct growth-promoting therapy for intestinal epithelium (Fig 1G-I). More importantly, in enteropathic mice, this approach accomplished complete histological reversal of disease pathology, restoring normal villus architecture and crypt depth (Fig 1G-I). This represents the first therapeutic application GSK3 inhibition as a WNT pathway agonist to successfully treat intestinal damage while enhancing normal tissue function— providing a facile application of selective pharmacology for intestinal regeneration.^47^ Importantly, while treated small intestines showed the expected physiological hyperplasia with enhanced villus height and crypt depth, comprehensive histological analysis revealed zero adenoma formation among 2,196 crypts examined across five mice (Figure S1F), confirming that WNT-mediated regeneration occurs without neoplastic transformation.

Colon-specific delivery confirms stem cell enhancement: We extended these findings to the mouse colon using targeted enema delivery of GSK3 inhibitor nanoparticles three times weekly for two weeks. We used LGR5-EGFP reporter mice to directly identify intestinal stem cells. ^48^ Immunohistochemistry for LGR5-GFP revealed increased expression in the crypt base of treated mice, with both an elevated number of GFP-positive cells per crypt (Fig. 1J,K) and increased crypt depth and overall tissue thickness (Fig. 1L), confirming that GSK3 inhibitor nanoparticle therapy induced augmented colon tissue and stem cell function. Parallel safety analysis in colon tissue demonstrated enhanced crypt depth consistent with stem cell activation, yet revealed zero adenomas or aberrant crypt foci among 1,664 crypts examined across three mice following enema delivery (Figure S1G), further validating that WNT hyperactivation promotes physiological regeneration without initiating tumorigenesis. To assess functional stem cell enhancement, we isolated primary colon crypts 24 hours after the final treatment and performed clonogenic assessment across a dose response of GSK3 inhibitor. Crypts from in vivo GSK3 inhibitor-treated mice demonstrated clonogenic potential and growth capacity even without in vitro GSK3 inhibitor or WNT agonist supplementation, while crypts from control mice exhibited no clonogenic capacity without exogenous GSK3 inhibitor (Fig. 1M). Remarkably, dose-response analysis revealed that the maximal growth potential of crypts from either in vivo treatment group was identical, indicating that GSK3 inhibition primes stem cells for enhanced responsiveness without fundamentally altering their final proliferative ceiling. This clonogenic enhancement persisted at least 24 hours post-treatment, demonstrating durable augmentation of stem cell competency. These findings establish that pharmacological GSK3 inhibition effectively activates WNT signaling in vivo, durably enhances intestinal stem cell function, and promotes regenerative growth across both small intestine and colon tissues in healthy and diseased states.

### OverWNTing colonic neoplasia

Over-WNTing" uncovers a critical vulnerability along the WNT signaling continuum. Conventional cancer therapy targets oncogenes through inhibition, yet we hypothesized that cancer cells might occupy a precarious position along the WNT signaling continuum where hyperactivation could prove therapeutically superior to suppression. To test this paradigm-challenging concept, we systematically evaluated WNT hyperactivation effects across the neoplastic spectrum using organoid lines with progressive mutational profiles reflecting typical colon cancer evolution: *Apc*^-/-^ (A); *Apc*^-/-^ *Kras*^G12D^, *Trp53*^-/-^ (AKP); and *Apc*^-/-^, *Kras*^G12D^, *Trp53*^-/-^ *Smad4*^-/-^ (AKPS). Strikingly, GSK3 inhibition demonstrated growth suppression of all neoplastic organoid lines regardless of their mutational complexity, with no concentration inducing growth enhancement. This contrasted sharply with normal mouse colon organoids, where GSK3 inhibition promoted dose-dependent growth (Fig 2A). This differential response reveals that while normal cells thrive with enhanced WNT signaling, cancer cells—despite their dependence on WNT—are killed by further pathway activation, a phenomenon we term "over-WNTing."

**Figure 2:**
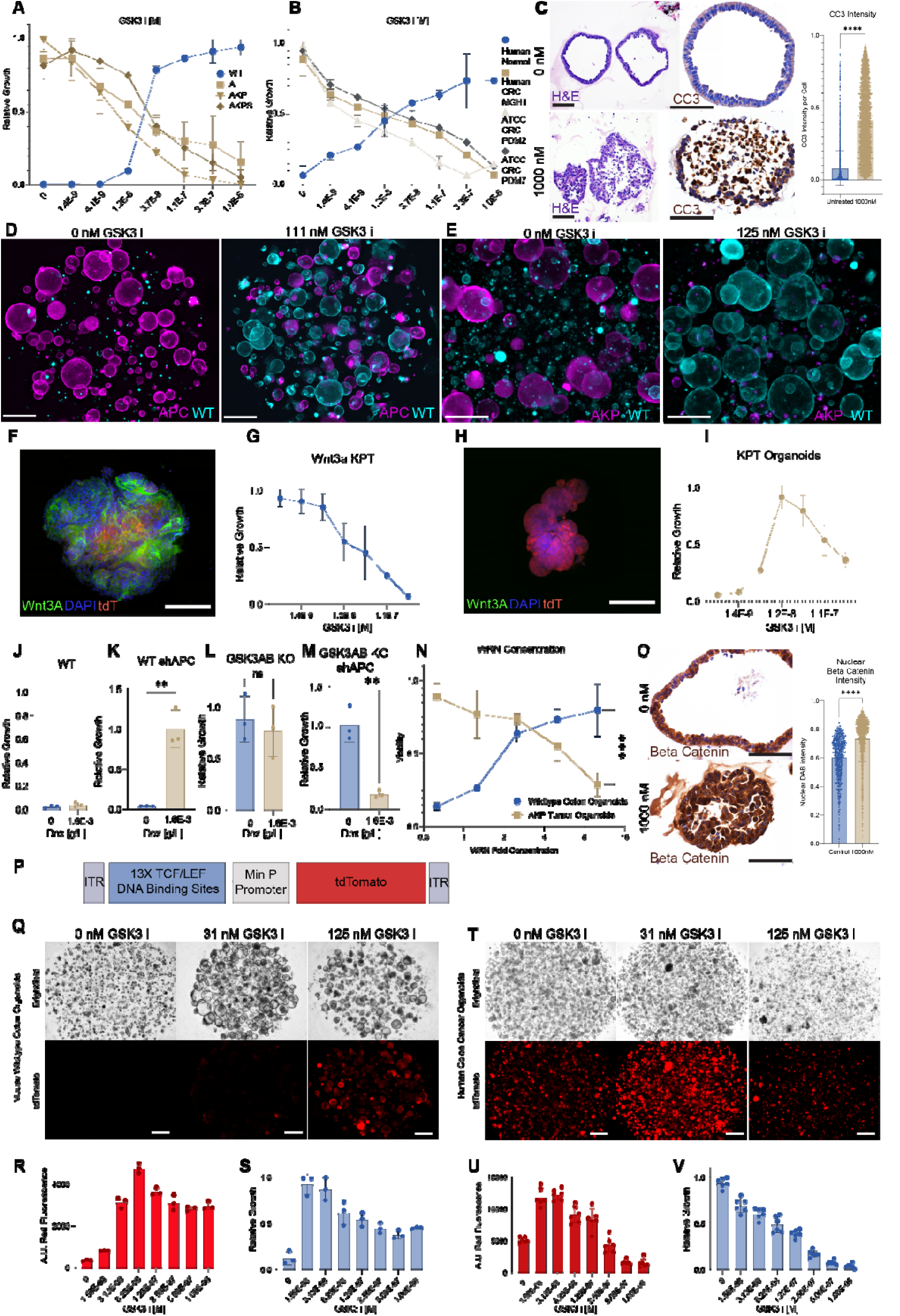
GSK3 inhibition inhibits colonic neoplasia through WNT signaling. (A) Dose-response curves of mouse colon organoid growth with GSK3i (LY2090314) treatment. Genotypes tested: wildtype (WT), Apc^-/-^ (A), Apc^-/-^;Kras^G12D^;Trp53^-/-^ (AKP), and Apc^-/-^;Kras^G12D^;Trp53^-/-^; Smad4^-/-^ (AKPS). Growth measured by resazurin assay after 72 hours and normalized to vehicle control for each genotype. n=3 biological replicates. (B) Dose-response curves of human colon organoid growth with GSK3i treatment. Normal colon organoids and patient-derived colorectal cancer organoids (CRC lines: MGH1, PDM2, PDM7) were cultured for 72 hours. Growth measured by resazurin assay and normalized to vehicle control. n=3 biological replicates. (C) Representative histological images of AKP mouse colon cancer organoids showing the effect of GSK3 inhibitor treatment. Top row: untreated control organoids (0 nM). Bottom row: organoids treated with 1000 nM GSK3 inhibitor (LY2090314) after 36 hours. Left panels show hematoxylin and eosin (H&E) staining demonstrating normal organoid architecture in control conditions versus disrupted morphology with cellular debris following GSK3 inhibitor treatment. Right panels show immunohistochemical staining for cleaved caspase-3 (CC3), a marker of apoptosis. Brown staining indicates CC3-positive apoptotic cells, which are markedly increased in the GSK3 inhibitor-treated organoids compared to untreated controls. Scale bars = 50 μm. Right: Quantification of cleaved caspase 3 intensity per cell (control: n = 3,797 cells from 13 fields; treated: n = 4,677 cells from 14 fields; 3 biological replicates each; ****p < 0.0001, unpaired t-test). (D,E) Competition assays between fluorescently labeled organoids cultured in media lacking WNT agonists. (D) Wildtype organoids (cyan) co-cultured with Apc^-/-^ organoids (magenta) ± GSK3i (111 nM). (E) Wildtype organoids (cyan) co-cultured with AKP organoids (magenta) ± GSK3i (125 nM). Images taken after 5 days of co-culture. (F-I) Validation of autocrine WNT3A expression effects. (F) Immunofluorescence of Kras^G12D^;Trp53^-/-^; tdTomato organoids with EF1A-driven Wnt3a transgene (Wnt3a-KPT). WNT3A (green), Hoechst 33342 (blue), tdTomato (red). (G) Growth curve of Wnt3a-KPT organoids with GSK3i dose response. (H) Control KPT organoids lacking Wnt3a transgene show no WNT3A staining. (I) Growth curve of control KPT organoids with GSK3i dose response. n=3 biological replicates for (G) and (I). (J-M) shRNA-mediated Apc knockdown experiments in WNT agonist-free media. Growth measured after 72 hours ± doxycycline (2 μg/ml) to induce shApc. (J,K) Wildtype organoids without (J) or with (K) shApc induction. (L,M) GSK3A/B double knockout organoids without (L) or with (M) shApc induction. n=3 biological replicates. (N) Dose-response curves of wildtype and AKP organoids cultured with concentrated WNT3A-conditioned media (WRN supernatant). Growth measured by resazurin assay after 72 hours. n=3 biological replicates. (O) Immunohistochemical staining for beta-catenin in mouse colon cancer organoids. Top panel: untreated control organoids (0 nM) showing baseline beta-catenin expression localized to cell membranes and partially to nuclei. Bottom panel: organoids treated with 1000 nM GSK3 inhibitor (LY2090314), 36 hours of treatment, demonstrating marked accumulation of beta-catenin with intense nuclear and cytoplasmic staining, consistent with hyperactivation of canonical WNT signaling. The increased beta-catenin levels correlate with the induction of apoptosis observed in these "over-WNTed" tumor organoids. Scale bars = 50 μm. Right: Quantification of nuclear beta-catenin intensity (control: n = 805 cells from 10 fields; treated: n = 1,608 cells from 13 fields; 3 biological replicates each; ****p < 0.0001, unpaired t-test). (P-V) WNT pathway activity reporter analysis. (P) Schematic of 13xTCF-tdTomato (TOP/tdTomato) reporter construct. (Q,R,S) Wildtype mouse organoids transduced with TOP/tdTomato reporter: (Q) representative brightfield and fluorescence images, (R) quantification of tdTomato fluorescence intensity per organoid, (S) corresponding growth measurements. (T,U,V) Human CRC organoids (PDM7) transfected with TOP/tdTomato reporter: (T) representative images, (U) quantification of tdTomato fluorescence intensity per organoid, (V) corresponding growth measurements. All measurements after 72 hours of GSK3i treatment. n=4 biological replicates. Scale bars: 500 μm (D,E,P,S); 100 μm (F,H) 50 μm (C). All growth assays measured by resazurin metabolic activity. GSK3i = LY2090314. Data shown as mean ± SD. Statistical analysis by unpaired two-tailed t-test (C) or one-way ANOVA with Tukey’s post-hoc test (other panels). ***p<0.001, **p<0.01.

**Figure S2:**
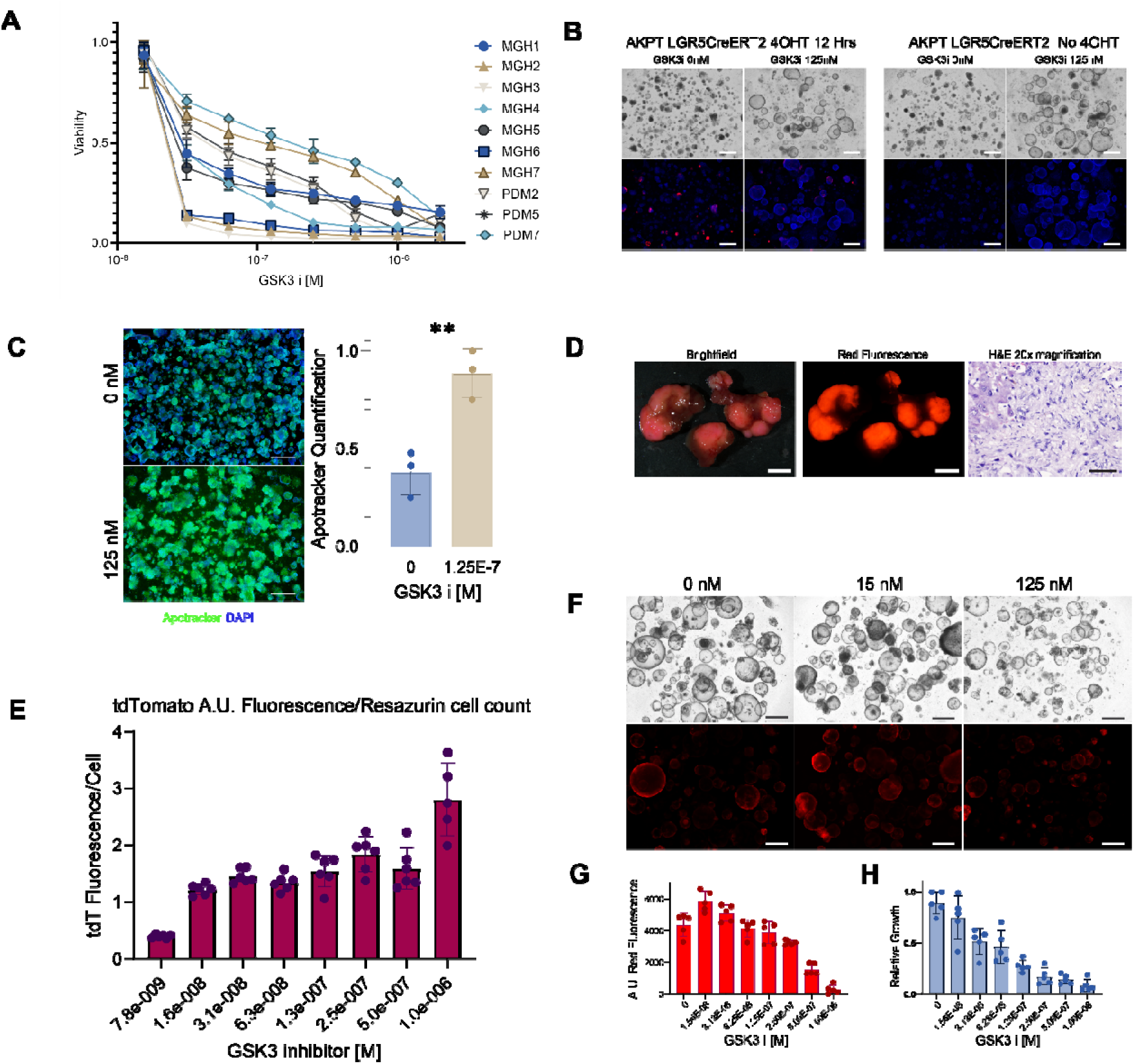
Human colon cancer organoid panel, effect of GSK3i on nascent tumorigenesis, TOP/tdTomato reporter validation in mouse colon cancer organoids, and Wnt3a-KPT organoid malignancy in vivo. (A) Dose-response curves of GSK3i (LY2090314) treatment across a panel of patient-derived human colorectal cancer organoids. Growth measured by resazurin metabolic assay after 72 hours, normalized to vehicle control. n=10 individual patient samples; demographic and clinical data provided in Supplementary Table 1. Data shown as mean ± SD. (B) Assessment of GSK3i effects on nascent tumor formation. Apc^fl/fl^;Kras^LSL-G12D^;Trp53^fl/fl^;Rosa26^LSL-tdTomato^;Lgr5-CreERT2 organoids were treated with 4-hydroxytamoxifen (500 ng/ml) for 12 hours to induce Cre-mediated recombination, then cultured ± GSK3i (100 nM) for 4 days. Red fluorescence indicates successful Cre recombination and tumor allele activation. DAPI (blue) marks all cell nuclei; DAPI-positive/tdTomato-negative cells represent untransformed organoids that escaped recombination. Representative images from n=3 independent experiments. (C) Apoptosis detection in AKP tumor organoids. Representative fluorescence images and quantification using Apotracker dye after 24 hours of GSK3i treatment (100 nM). n=4 biological replicates. **p<0.01, unpaired t-test. (D) In vivo tumorigenicity of Wnt3a-KPT organoids. Left: Gross brightfield and fluorescence images of mouse liver 5 weeks after orthotopic implantation of Kras^G12D^;Trp53^-/-^;tdTomato organoids expressing EF1A-driven Wnt3a transgene (Wnt3a-KPT). Right: Hematoxylin and eosin staining confirming tumor formation. Representative images from n=5 mice. (E) Normalized red fluorescence of TOP/tdTomato reporter in human CRC organoids, normalized by cell number, primary data found in Figure 2 U,V. demonstrating WNT reporter activity inversely correlating with cell number across all concentrations. (F,G,H) WNT pathway reporter analysis in mouse colorectal cancer organoids. (F) Representative brightfield (top) and tdTomato fluorescence (bottom) images of Apc^-/-^;Kras^G12D^;Trp53^-/-^ (AKP) organoids transduced with 13xTCF-tdTomato (TOP/tdTomato) reporter, cultured with GSK3i dose response for 72 hours. (G) Quantification of tdTomato fluorescence intensity per well from (F). (H) Corresponding growth measurements of AKP organoids by resazurin assay. n=5 biological replicates for (G) and (H). Data shown as mean ± SD. GSK3i = LY2090314 for all experiments. Scale bars: 500 μm (B,C,F); 5 mm (D, gross images); 200 μm (D, histology).

Human cancer organoids validate the therapeutic paradigm. We extended these findings to human colorectal cancer using organoid lines derived from seven patients with diverse clinical characteristics spanning multiple disease stages and molecular subtypes (Table 1), plus three commercially available human colon cancer lines (Fig. S2A). The results precisely mirrored our mouse studies: GSK3 inhibition consistently suppressed all human CRC organoids while promoting normal human organoid growth (Fig 2B), demonstrating that "over-WNTing" represents a conserved therapeutic vulnerability across species and genetic backgrounds. Mechanistic investigation revealed that GSK3 inhibition induced apoptosis in tumor organoids, as demonstrated by cleaved caspase-3 immunohistochemistry in mouse adenocarcinoma organoids (Fig. 2C). Following GSK3 inhibitor treatment for 36 hours, AKP organoids displayed extensive apoptotic cell death characterized by abundant cleaved caspase-3-positive cells throughout the disrupted organoid architecture. The treated organoids showed loss of normal epithelial organization with cellular debris and fragmentation, contrasting sharply with the intact columnar epithelium and minimal baseline apoptosis observed in untreated controls. Quantification across thousands of cells confirmed this profound apoptotic response, with virtually no cleaved caspase-3 signal in control organoids while GSK3 inhibitor treatment induced massive apoptosis (p < 0.0001, Fig. 2C). This widespread activation of the caspase-3 apoptotic pathway was further validated using Apotracker staining, which detects early apoptotic events through calcium-independent phosphatidylserine outer membrane exposure (Fig. S2C).

**Table 1:**
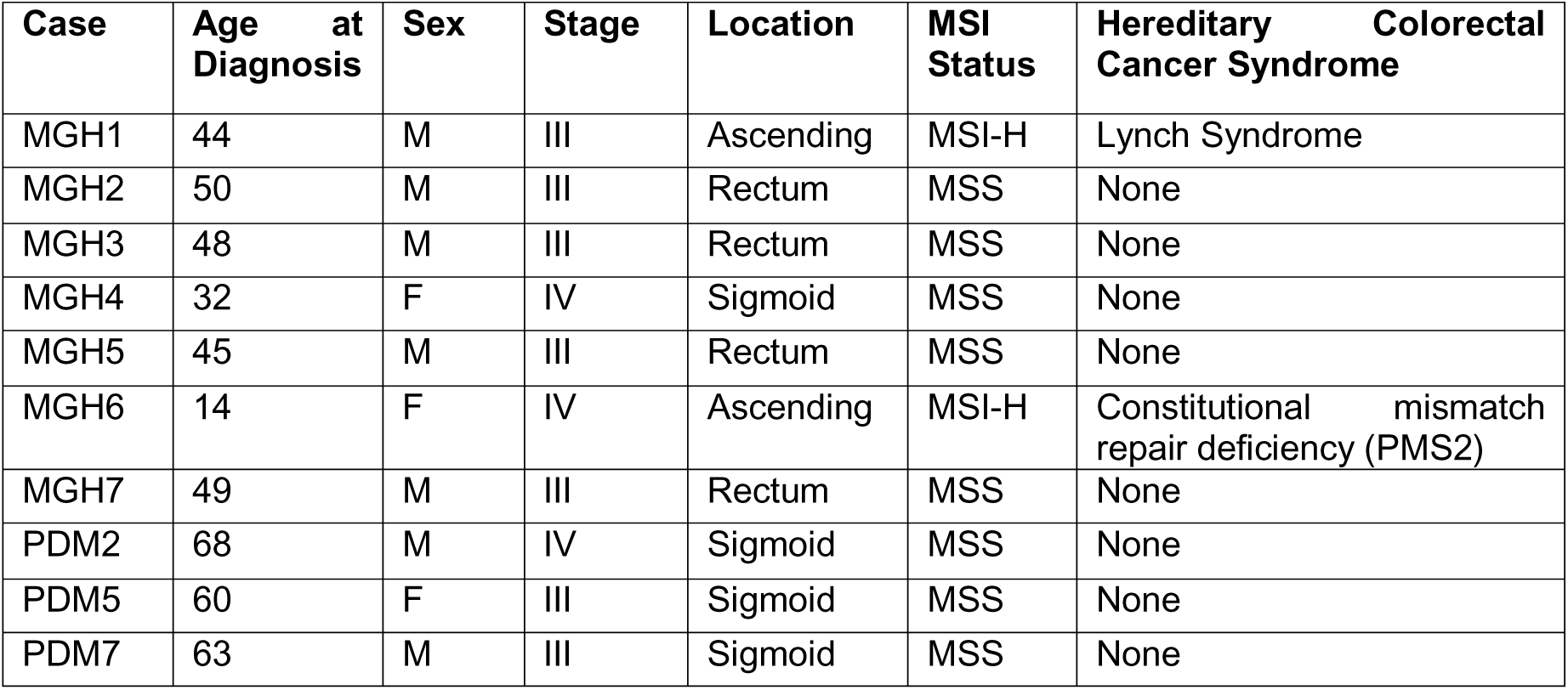
Demographic characteristics of patients from whom organoid lines were generated.

| Case | Age at Diagnosis | Sex | Stage | Location | MSI Status | Hereditary Colorectal Cancer Syndrome |
| --- | --- | --- | --- | --- | --- | --- |
| MGH1 | 44 | M | III | Ascending | MSI-H | Lynch Syndrome |
| MGH2 | 50 | M | III | Rectum | MSS | None |
| MGH3 | 48 | M | III | Rectum | MSS | None |
| MGH4 | 32 | F | IV | Sigmoid | MSS | None |
| MGH5 | 45 | M | III | Rectum | MSS | None |
| MGH6 | 14 | F | IV | Ascending | MSI-H | Constitutional mismatch repair deficiency (PMS2) |
| MGH7 | 49 | M | III | Rectum | MSS | None |
| PDM2 | 68 | M | IV | Sigmoid | MSS | None |
| PDM5 | 60 | F | III | Sigmoid | MSS | None |
| PDM7 | 63 | M | III | Sigmoid | MSS | None |

Direct co-culture experiments dramatically illustrated this differential effect: without GSK3 inhibition, adenomatous organoids (magenta) grew robustly while normal organoids (cyan) failed to grow due to the absence of exogenous WNT agonists. GSK3 inhibition completely reversed this pattern, supporting normal organoid growth while inhibiting tumor organoids (Fig 2D). This effect extended to co-cultures of adenocarcinoma organoids (AKP) with normal organoids (Fig 2E). These competition experiments effectively demonstrate the selective therapeutic principle—normal and tumor cells respond oppositely to the same WNT-activating intervention. To examine the effect of GSK3 inhibition on tumor initiation, we developed a competition assay using *Apc*^fl/fl^ *LSL-Kras*^G12D^, *Trp53*^fl/fl^, Rosa-26 LSL-tdTomato; LGR5-CreERT2 organoids. Using low-dose 4-hydroxytamoxifen exposure to partially activate tumor alleles in a subset of LGR5+ stem cells, we created direct competition between nascent tumor clones and normal cells within the same organoid culture - modeling the early stages of tumor initiation where only rare stem cell-initiated clones acquire transforming mutations. Control cultures readily established red fluorescent tumor clones that persisted and expanded, however, GSK3 inhibition following 4-hydroxytamoxifen exposure completely eliminated these nascent tumor clones within 4 days, despite their genetic activation, instead selecting for the growth of normal organoids (blue) (Fig. S2B). This immediate elimination of nascent tumor clones reveals a fundamental safety mechanism inherent to WNT agonism therapy. While GSK3 inhibition enhances WNT signaling in normal cells to promote regeneration, any cell that undergoes APC loss—the irreversible genetic switch defining colorectal cancer—immediately experiences lethal over-WNTing. This occurs because APC loss is a binary genetic commitment that permanently locks cells into a constitutively high WNT state. The moment a normal stem cell loses APC through sporadic mutation, it jumps from the safe physiological WNT range where GSK3 inhibition or other WNT agonist is beneficial, to the precarious tumor setpoint where additional WNT signaling becomes lethal. This creates continuous surveillance: GSK3 inhibitor therapy simultaneously enhances normal tissue function while automatically eliminating any cell that undergoes transformation. It is this expanded concept of specific setpoints along the WNT signaling continuum that leverages the step-function nature of APC loss to provide intrinsic selectivity—the genetic event that initiates cancer also triggers its immediate destruction under treatment conditions. This transforms the theoretical liability of WNT enhancement into a therapeutic asset, as the treatment can not cause adenomas because it eliminates transformed cells immediately as they meet the definition, genetic APC loss.

To dissect whether over-WNTing represents a fundamental vulnerability of high WNT states rather than an APC-specific synthetic lethality, we engineered an orthogonal model of constitutive WNT activation. We expressed transgenic WNT3a in *Kras*^G12D^; *Trp53*^-/-^; TdTomato (KPT) organoids, creating tumor cells that achieve niche independence through autonomous WNT ligand production rather than destruction complex disruption. Immunofluorescence confirmed successful engineering, revealing membranous WNT3a localization in transgenic organoids while parental KPT controls showed no detectable signal (Fig. 2F,H). Notably, these WNT3a-expressing tumor organoids recapitulated the over-WNTing vulnerability observed in APC-mutant cells. GSK3 inhibition induced dose-dependent growth suppression in WNT3a KPT organoids, mirroring the response of *Apc*^-/-^ organoids despite achieving constitutive WNT activation through an entirely different mechanism (Fig. 2G). This convergent phenotype establishes that susceptibility to over-WNTing arises from the high WNT state itself, not from specific disruption of the destruction complex, and not only through a specific interaction between APC loss and GSK3 inhibition.

The parental KPT organoids provided additional mechanistic insight, exhibiting a striking biphasic response to GSK3 inhibition that directly demonstrates the WNT-just-right principle. Low concentrations of GSK3 inhibitor rescued their exogenous WNT dependence and enabled growth, while higher concentrations pushed them beyond their optimal signaling threshold and induced growth arrest (Fig. 2I). This dose-response curve captures the precarious balance cancer cells must maintain—sufficient WNT signaling for survival, but not so much as to trigger cell death. The WNT3a KPT organoids retain full tumorigenic potential, forming lethal tumors upon transplantation that phenocopy AKP adenocarcinomas (Fig. S2D). This demonstrates that these CRC organoids actually maintain WNT3a overexpression as a mechanism of niche independence, similar to APC loss, but even an orthogonal method of niche independence can lead to overWNTing vulnerability.

To determine whether the incompatibility between APC loss and GSK3 inhibition represents true synthetic lethality, we tested the reciprocal interaction: whether APC knockdown would inhibit growth in GSK3-deficient cells. We generated organoids with doxycycline-inducible sh*Apc* to temporally control APC expression in different genetic backgrounds. In normal organoids, doxycycline-induced APC knockdown promoted WNT-independent growth as expected (Fig. 2K), while doxycycline alone had no effect on untransduced controls (Fig. 2J). GSK3A/B double knockout organoids similarly showed no response to doxycycline alone (Fig. 2L). However, when we induced APC knockdown in GSK3A/B knockout organoids, we observed growth inhibition rather than the enhanced growth seen in normal organoids (Fig. 2M). This reciprocal effect demonstrates that simultaneous functional loss of APC and GSK3—regardless of whether achieved through pharmacologic inhibition, genetic deletion, or knockdown—creates a non-viable cellular state. The synthetic lethality is bidirectional: GSK3 inhibition kills APC-mutant cells, and APC loss kills GSK3-deficient cells. Critically, achieving the same growth inhibition through purely genetic means i.e. GSK3A/B knockout with sh*APC*, definitively establishes that this synthetic lethality reflects a fundamental liability in WNT high niche independent states, and not a specific interaction due to pharmacologic GSK3 inhibition. These findings reveal that CRC cells have an absolute ceiling for tolerable WNT pathway activation, beyond which further signaling increases become lethal.

To test whether WNT hyperactivation vulnerability is intrinsic to high-WNT states rather than specific to GSK3 manipulation, we developed a novel protocol to concentrate native WNT3A, R-Spondin, and Noggin proteins. Previous approaches require cytotoxic detergents for protein solubilization, ^49,50^ while newer bispecific WNT agonists exhibit hook effects that paradoxically reduce activity at high concentrations. Our detergent-free method achieved 20-fold protein concentration while preserving bioactivity. Concentrated WRN preparations replicated the exact differential growth pattern: dose-dependent tumor organoid inhibition with simultaneous normal organoid enhancement (Fig. 2N). This pure ligand approach demonstrates that ’over-WNTing’ is not only pathway conceptual but actually literal: excess WNT proteins themselves are sufficient to kill cancer cells while enhancing normal cell growth. Together with our genetic experiments, this establishes that niche independent cancer cells cannot tolerate further pathway activation regardless if achieved through GSK3 inhibition, genetic disruption, or physiological ligand stimulation.

To directly visualize the impact of GSK3 inhibition on WNT pathway activation at the cellular level, we performed beta-catenin immunohistochemistry on AKP tumor organoids. In untreated controls, beta-catenin showed the expected pattern for APC-deficient cells—prominent membranous staining with some nuclear localization reflecting constitutive WNT activation (Fig. 2O). GSK3 inhibitor treatment dramatically amplified this signal, inducing total beta-catenin accumulation throughout the organoids, with some demonstrating primarily nuclear beta-catenin, demonstrating that WNT signaling can be pushed beyond the constitutive activation imposed by APC loss. Quantification across thousands of cells confirmed this hyperactivation, with nuclear beta-catenin intensity increasing significantly in treated organoids (p < 0.0001, Fig. 2O). Notably, the cells displaying the most intense nuclear beta-catenin staining corresponded morphologically to the apoptotic cells identified by cleaved caspase-3 immunostaining (Fig. 2C), with disrupted cellular architecture and condensed nuclei. This correlation between increased nuclear beta-catenin accumulation and apoptotic morphology provides direct evidence that pushing WNT signaling beyond its already elevated state in APC-mutant tumors triggers cell death. The transition from constitutive to hyperactivated WNT signaling—manifested as increased total beta-catenin with enhanced nuclear localization—represents the molecular threshold where oncogenic signaling becomes lethal "over-WNTing."

To quantitatively measure WNT pathway dynamics in living organoids and directly test whether APC-mutant cells retain capacity for further pathway activation, we engineered a high-sensitivity fluorescent reporter system. We constructed a synthetic promoter containing 13 tandem beta-catenin/TCF consensus sites (AGATCAAAGG) driving tdTomato expression, integrated via PiggyBac transposition for stable expression across passages (Fig. 2P). This design provides superior sensitivity compared to traditional 7-8x TCF reporters and enabled continuous measurements in live 3D organoids—capabilities not achievable with standard luciferase-based assays. ^51^ The reporter faithfully tracked WNT signaling dynamics in normal colon organoids. GSK3 inhibition induced dose-dependent reporter activation that precisely correlated with normal organoid growth at physiological concentrations (Fig. 2Q-S). Notably, both growth and reporter fluorescence plateaued at higher GSK3 inhibitor concentrations within the on-target range, demonstrating that normal colon cells can buffer WNT pathway hyperactivation.

When introduced into human colon adenocarcinoma organoids, the reporter demonstrated strong baseline activity without GSK3 inhibition, consistent with APC mutation-driven constitutive WNT activation. Critically, GSK3 inhibition further increased WNT reporter activity in a dose-dependent manner (Fig. 2T,U), definitively establishing that APC-mutant cells are not at maximal WNT signaling. Growth analysis of these same organoids revealed the therapeutic paradox: while untreated cancer organoids grew robustly, increasing GSK3 inhibitor concentrations that enhanced WNT reporter activity proved progressively toxic (Fig. 2V). When normalized to cell number, the data revealed a clear linear relationship—increased per-cell WNT signaling directly correlated with decreased viability (Fig. S2E). Mouse colon cancer organoids displayed similar patterns when expressing the reporter (Fig. S2 F,G,H). These findings provide direct molecular evidence of over-WNTing: APC-mutant cancer cells exist at a precarious WNT optimum rather than maximal activation. Further pathway stimulation—whether through GSK3 inhibition, genetic manipulation, or excess WNT proteins—pushes these cells beyond tolerable signaling thresholds. This creates an unexpected therapeutic vulnerability where the very pathway driving cancer growth becomes lethal when hyperactivated.

### Wnt hyperactivation effectively treats colon cancer *in vivo*

Having established that WNT hyperactivation selectively kills cancer cells while enhancing normal tissue, in vitro, we tested whether this therapeutic window could be exploited in vivo. We selected peritoneal carcinomatosis as our primary model—a devastating manifestation of advanced colorectal cancer that remains essentially untreatable and carries median survival of only 6-12 months.^52^ This model stringently tests both efficacy and safety, as intraperitoneally delivered therapy directly contacts both tumors and normal intestinal epithelium. We established disseminated disease by injecting fluorescent AKPT colon cancer organoids throughout the peritoneal cavity, allowing three days for tumor establishment before initiating daily GSK3 inhibitor nanoparticle therapy (Fig. 3A). The results exceeded conventional therapeutic benchmarks: while control mice developed extensive tumor burden throughout the peritoneum, one week of GSK3 inhibitor nanoparticle treatment achieved 93% reduction in total tumor burden—near-complete disease eradication (Fig. 3B,C). Individual tumor counts decreased from 15-20 per animal to fewer than 5, with remaining lesions showing minimal fluorescence.

**Figure 3:**
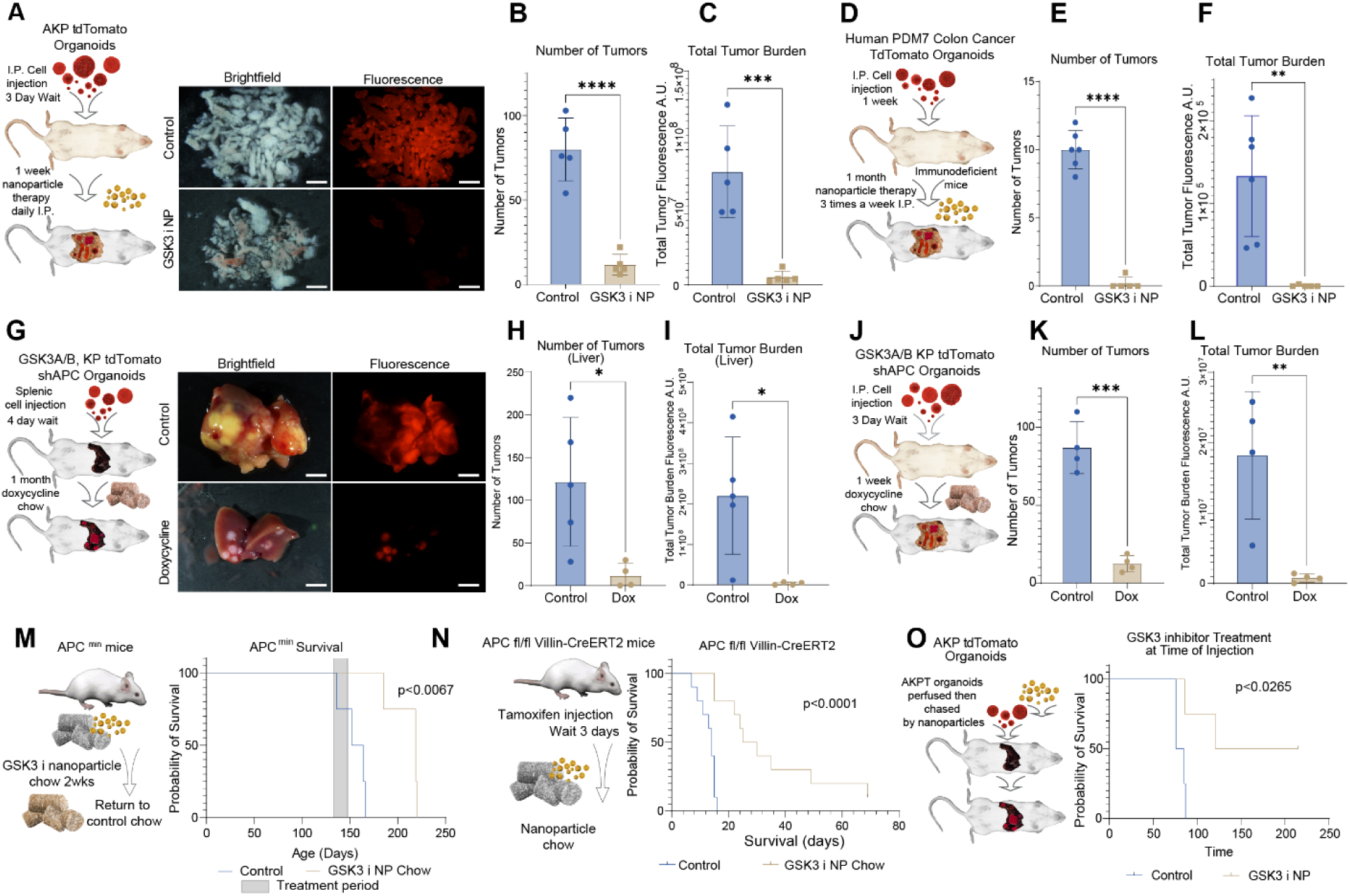
Wnt hyperactivation effectively treats colon cancer *in vivo*. (A-C) Peritoneal carcinomatosis model using mouse colorectal cancer organoids. (A) Experimental schematic: AKP-tdTomato organoids injected intraperitoneally, followed by daily treatment with control or GSK3i-loaded PLGA nanoparticles starting at day 3 for 7 days. Representative gross brightfield and fluorescence images of peritoneal tumor burden at endpoint. n=5 mice per group, representative of 3 independent experiments. (B) Quantification of tumor nodule count per mouse. (C) Quantification of total tumor burden calculated as Σ(mean fluorescence intensity × tumor area) for all tumors per mouse. ***p<0.001, unpaired t-test. (D-F) Peritoneal carcinomatosis model using patient-derived human colorectal cancer organoids. (D) Experimental schematic: human CRC organoids injected intraperitoneally into NSG mice, followed by treatment with control or GSK3i nanoparticles three times weekly starting at day 7 for 4 weeks. (E) Quantification of tumor nodule count per mouse. (F) Quantification of total tumor burden. n=6 mice per group, representative of 2 independent experiments. **p<0.01, unpaired t-test. (G-I) Liver metastasis model with inducible Apc knockdown. (G) Experimental schematic: GSK3A/B knockout KPT organoids carrying doxycycline-inducible shApc were implanted via splenic injection. Mice received control or doxycycline chow starting at day 3. Representative images at 4-week endpoint. n=5 mice per group, representative of 3 independent experiments. (H) Quantification of liver tumor nodule count. (I) Quantification of total liver tumor burden. ****p<0.0001, unpaired t-test. (J-L) Peritoneal carcinomatosis model with inducible Apc knockdown. (J) Experimental schematic: GSK3A/B knockout KPT shApc organoids injected intraperitoneally, followed by control or doxycycline chow starting at day 3. (K) Quantification of tumor nodule count at 3-week endpoint. (L) Quantification of total tumor burden. n=4 mice per group. *p<0.05, unpaired t-test. (M) Survival analysis of Apc^Min/+^ mice. Mice received GSK3i nanoparticle-supplemented chow for 2 weeks followed by return to normal chow. Survival monitored up to 300 days. n=4 mice per group, representative of 2 independent experiments. Log-rank test. (N) Survival analysis following acute Apc deletion. Apc^fl/fl^;Villin-CreERT2 mice received tamoxifen (i.p.) followed by control or GSK3i-supplemented chow starting at day 3. n=10 mice per group, representative of 3 independent experiments. Log-rank test. (O) Survival analysis following metastatic tumor challenge. AKP organoids implanted via splenic injection with immediate treatment using control or GSK3i nanoparticles. n=4 mice per group, representative of 2 independent experiments. Log-rank test. GSK3i = LY2090314 for all experiments. Scale bar: 5 mm. Data shown as mean ± SD for (B,C,E,F,H,I,K,L). Statistical analysis by unpaired two-tailed t-test or log-rank test as indicated. *p<0.05, **p<0.01, ***p<0.001, ****p<0.0001.

**Figure S3:**
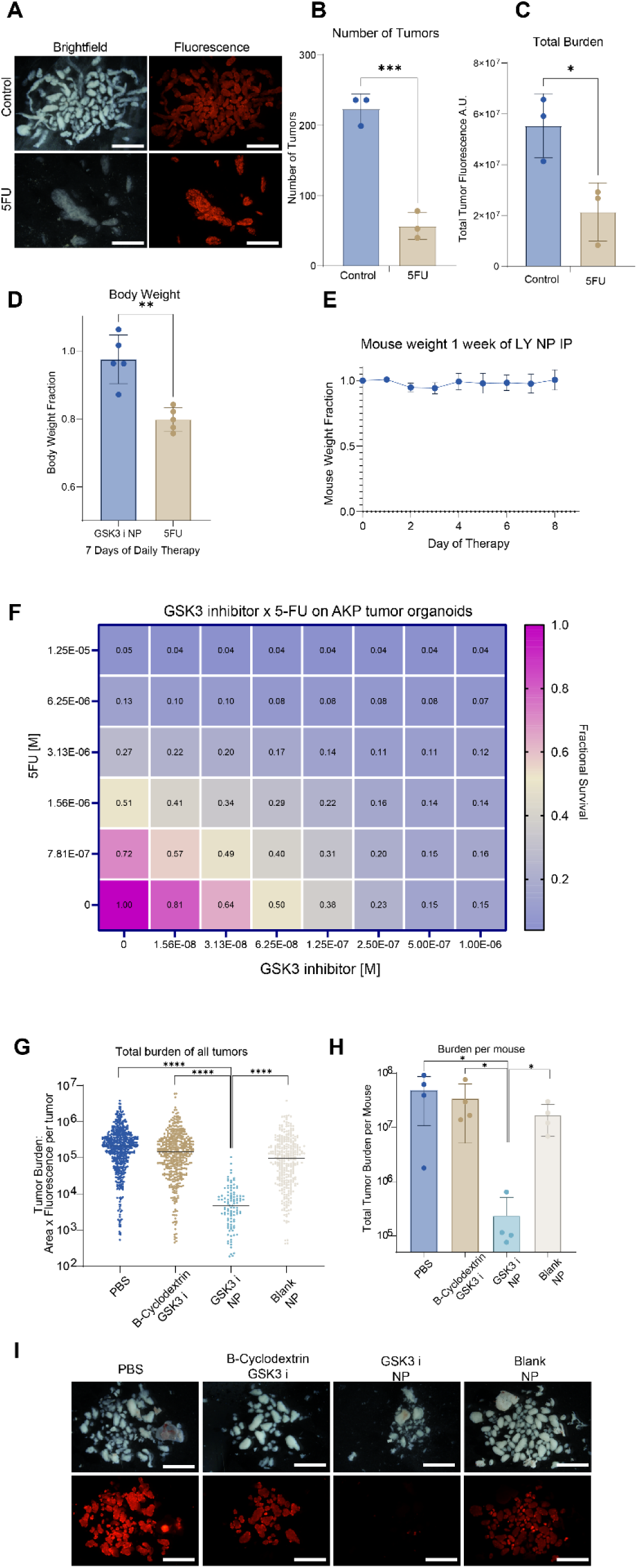
Comparison of 5-FU and GSK3i nanoparticle therapy in peritoneal carcinomatosis, and evaluation of GSK3i delivery methods. (A-C) 5-Fluorouracil treatment of peritoneal carcinomatosis. (A) Representative brightfield and fluorescence images of AKPT-tdTomato peritoneal tumors following 7-day treatment with vehicle control or 5-FU (50 mg/kg/day, i.p.). (B) Quantification of tumor nodule count per mouse. (C) Quantification of total tumor burden calculated as Σ(mean fluorescence intensity × tumor area). n=3 mice per group, representative of 3 independent experiments. Data shown as mean ± SD, unpaired t-test. (D) Body weight changes during therapy. Comparison of daily weights (normalized to initial weight) between GSK3i nanoparticle-treated mice (500 μl i.p. daily) and 5-FU-treated mice (50 mg/kg/day i.p.) over 7 days. n=5 mice per group, representative of 3 independent experiments. ****p<0.0001, two-way ANOVA. (E) Tolerability of GSK3i nanoparticle therapy. Daily weights of mice receiving GSK3i nanoparticles (500 μl i.p.) normalized to starting weight over 8-day treatment period. n=5 mice. Data shown as mean ± SD. (F) In vitro drug combination analysis. Dose-response matrix of GSK3i (LY2090314) combined with 5-FU on AKP mouse colorectal cancer organoids. Fractional survival normalized to untreated control. Heat map indicates synergy (blue) or antagonism (red). n=3 biological replicates. (G-I) Comparison of GSK3i delivery methods in peritoneal carcinomatosis model. AKPT organoids treated daily for 7 days (500 μl i.p.) with: PBS (vehicle control), β-cyclodextrin-solubilized GSK3i, GSK3i-loaded PLGA nanoparticles, or blank PLGA nanoparticles. β-cyclodextrin solution (20% w/v) contained equivalent LY2090314 dose as nanoparticle formulation. (G) Individual tumor burden measurements shown as scatter plot. (H) Total tumor burden per mouse. n=4 mice per group, representative of 2 independent experiments. *p<0.05, ****p<0.0001, one-way ANOVA with Tukey’s post-hoc test. (I) Representative gross brightfield and fluorescence images for each treatment condition. GSK3i = LY2090314 for all experiments. Scale bar: 1 cm. Data shown as mean ± SD unless otherwise noted.

To benchmark our approach against current clinical standards, we compared GSK3 inhibitor nanoparticles to 5-fluorouracil, the cornerstone chemotherapy for colorectal cancer. We administered 5-FU at its maximum tolerated dose (50 mg/kg daily)—the highest achievable without lethal toxicity, after determining that lower doses showed proportionally reduced efficacy. ^53^ This head-to-head comparison revealed the transformative potential of pathway hyperactivation over conventional cytotoxic therapy. While 5-FU significantly reduced tumor burden, achieving 63% reduction after one week of daily treatment, it fell substantially short of the 93% reduction achieved with GSK3 inhibitor nanoparticles (Fig. S3A-C). More striking was the toxicity differential: 5-FU caused severe systemic toxicity with 20% body weight loss, lethargy, and rough coat appearance—classic markers of chemotherapy-induced morbidity (Fig. S3D). In stark contrast, GSK3 inhibitor nanoparticle-treated mice maintained stable weight and normal activity throughout treatment, appearing indistinguishable from healthy controls (Fig. S3E). This inverse relationship—superior efficacy with absent toxicity—exemplifies how enhancing rather than inhibiting driver pathways can overcome fundamental limitations of conventional cancer therapy. The therapeutic superiority depended critically on nanoparticle formulation. Neither beta-cyclodextrin-solubilized GSK3 inhibitor nor blank nanoparticles reduced tumor burden, despite delivering equivalent drug doses (Fig. S3G-I). Intriguingly, combining low-dose GSK3 inhibition with 5-FU showed additive effects in organoid studies (Fig. S3F), suggesting our approach could either replace maximum-dose chemotherapy or enable reduced-toxicity combination regimens.

To address species-specific concerns and validate therapeutic efficacy against human cancer, we tested GSK3 inhibitor nanoparticles in patient-derived xenografts. We engrafted tdTomato-expressing human colorectal cancer organoids (PDM7) into immunodeficient mouse peritoneal cavities, allowing one week for tumor establishment to model clinically relevant disease burden before initiating therapy (Fig. 3D). This extended treatment protocol—one month of thrice-weekly nanoparticle administration—tested both sustained efficacy and potential resistance development. Human tumors proved equally susceptible to WNT hyperactivation as mouse tumors. GSK3 inhibitor nanoparticle treatment reduced human tumor counts by over 80% and achieved near-complete elimination of total tumor burden (Fig. 3E,F). Remarkably, this efficacy was maintained throughout the month-long treatment period without evidence of resistance emergence or reduced responsiveness. The sustained response of patient-derived tumors confirms that vulnerability to over-WNTing is conserved across species and validates the translational potential of targeting high-WNT states in human colorectal cancer.

While nanoparticle delivery demonstrated therapeutic efficacy, establishing that over-WNTing reflects fundamental biology rather than pharmacological artifact required genetic validation in vivo. We engineered an inducible synthetic lethality system using GSK3A/B knockout KPT organoids carrying doxycycline-regulated shApc—enabling temporal control of the precise genetic interaction that causes tumor inhibition in vitro. This orthogonal approach eliminates all concerns about drug specificity, off-target effects, or pharmacokinetic limitations. We tested this genetic over-WNTing in the most clinically relevant metastatic site—the liver, where 70% of colorectal cancer patients develop metastases.^54^ Following splenic injection and surgical splenectomy, we allowed four days for micrometastatic seeding before initiating doxycycline treatment. Genetic induction of the synthetic lethal state potently suppressed metastatic outgrowth: while control mice developed extensive liver metastases, doxycycline-treated mice showed >75% reduction in both tumor number and total metastatic burden after one month (Fig. 3G-I). This genetic approach proved equally effective against established peritoneal carcinomatosis. One week of doxycycline-induced APC knockdown—triggering over-WNTing in GSK3-null tumors—replicated the therapeutic efficacy of GSK3 inhibitor nanoparticles (Fig. 3J-L).

The convergence of pharmacological and genetic approaches on identical therapeutic outcomes provides definitive proof that vulnerability to WNT hyperactivation is an intrinsic property of high-WNT cancer cells, exploitable through multiple independent mechanisms. Achieving tumor regression in preclinical models of CRC, through pure genetic interaction—without any small molecule involvement— establishes that this phenomenon reflects fundamental cancer cell biology rather than pharmacological artifacts. This genetic validation demonstrates a new therapeutic principle: the very pathways driving cancer can be leveraged against tumors when pushed beyond tolerable thresholds.

To evaluate therapeutic potential across the adenoma-carcinoma sequence, we tested GSK3 inhibitor nanoparticles in autochthonous models that recapitulate distinct clinical scenarios. In Apc^min^ mice—the gold standard model for familial adenomatous polyposis—two weeks of oral GSK3 inhibitor nanoparticle administration significantly extended survival (Fig. 3M). ^55^ We confirmed this preventive potential using temporally controlled tumor initiation. In Apc^fl/fl^ Villin-CreERT2 mice, initiating GSK3 inhibitor nanoparticle treatment just three days after tamoxifen-induced Apc loss—capturing the earliest moments of cellular transformation—significantly extended survival compared to untreated controls (Fig. 3N).

Most clinically relevant for patients with advanced stage colorectal cancer, we modeled post-surgical adjuvant therapy by administering a single GSK3 inhibitor nanoparticle dose immediately after splenic injection of AKPT organoids. Remarkably, this one-time intervention—delivered when micrometastases are seeding the liver—significantly extended survival (Fig. 3O), and some mice never developed metastases. Since the GSK3 inhibitor nanoparticle therapy exhibits no observed toxicity while simultaneously enhancing normal intestinal function, it may be uniquely useful in the adjuvant setting to limit metastatic seeding.

### WNT hyperactivation induces RHOC-mediated apoptosis via non-canonical signaling

To elucidate the molecular mechanism underlying "over-WNTing"-induced tumor cell death, we performed comprehensive bulk RNA-seq transcriptomic analysis on AKP tumor organoids treated with GSK3 inhibitor concentrations spanning from sub-inhibitory to highly inhibitory doses (Fig. 4A). This dose-response design allowed us to capture the transition from tolerated WNT activation to lethal hyperactivation. Principal component analysis revealed that the primary transcriptomic axis (PC1) directly correlated with increasing GSK3 inhibition intensity. While sub-inhibitory doses (≤10 nM) that permitted organoid survival clustered with untreated controls, the growth-inhibitory concentrations (100-1000 nM) established distinct, reproducible gene expression signatures that progressed along PC1 in a dose-dependent manner (Fig. 4B). We performed hierarchical clustering of the 100 most differentially expressed genes to gain insight into the molecular drivers of "over-WNTing,". This unbiased analysis revealed RHOC—a member of the Rho GTPase family that interfaces with non-canonical WNT/planar cell polarity signaling—as prominently upregulated in the lethal WNT hyperactivation state (Fig. 4B, red box). DESeq2 differential expression analysis ^56^ confirmed *RHOC* as the 14th most significantly upregulated gene among 328 differentially expressed genes (adjusted p<0.01, Log2 fold change = 2.71) when comparing 1000 nM GSK3 inhibitor treatment to control conditions. *RHOC* expression demonstrated clear and specific dose-dependency, with minimal induction at sub-lethal concentrations but dramatic upregulation precisely correlating with the onset of tumor cell death (Fig. 4C).

**Figure 4:**
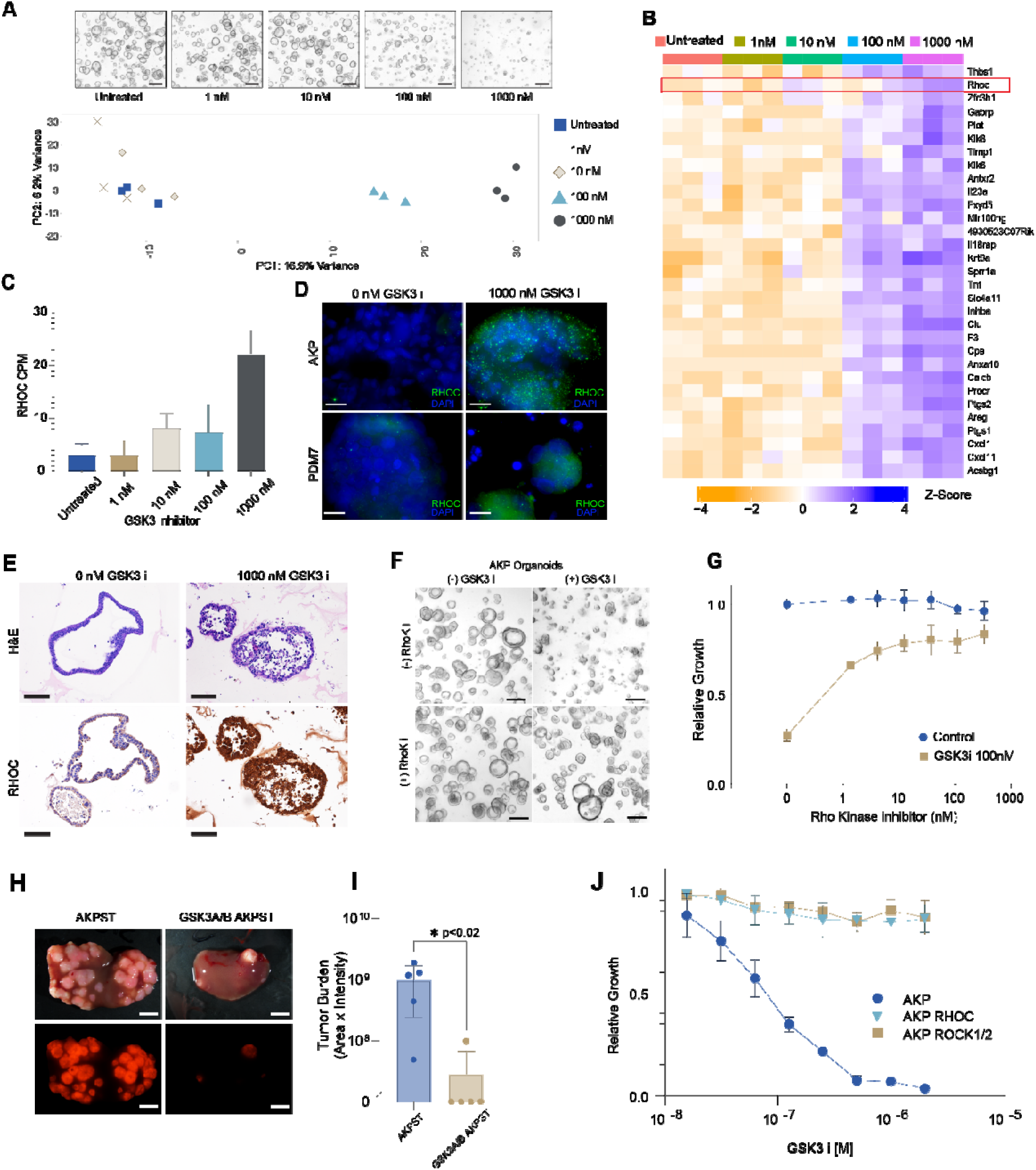
WNT hyperactivation induces RHOC-mediated apoptosis via non-canonical signaling. (A) Principal component analysis of bulk RNA-sequencing from AKPT organoids cultured with GSK3i dose response (0, 10, 100, 1000 nM LY2090314) for 72 hours. Representative brightfield images of organoids at each concentration shown below PCA plot. n=3 biological replicates per condition. Scale bar: 200 μm. (B) Unsupervised hierarchical clustering heatmap of differentially expressed genes from RNA-seq dataset in (A). Red box highlights *Rhoc* expression pattern across GSK3i concentrations. Color scale represents z-score of log2-transformed expression values. (C) *Rhoc* gene expression quantification across GSK3i doses. Data shown as counts per million mapped reads (CPM) from RNA-seq analysis. n=3 biological replicates, mean ± SD. One-way ANOVA with Tukey’s post-hoc test. (D) Immunofluorescence staining for RHOC protein (green) with DAPI nuclear counterstain (blue). Top: AKPT mouse organoids. Bottom: Patient-derived human colorectal cancer organoids (PDM7). Left: vehicle control. Right: 1000 nM GSK3i treatment for 72 hours. Representative images from n=3 independent experiments. Scale bar: 20 μm. (E) Representative histological images of AKP mouse colon cancer organoids following 36-hour treatment with GSK3 inhibitor after 2 days of growth. Top panels: Hematoxylin and eosin (H&E) staining. Bottom panels: RHOC immunohistochemical staining. Left column: untreated control organoids (0 nM GSK3 i) showing normal organoid architecture with minimal RHOC expression. Right column: organoids treated with 1000 nM GSK3 inhibitor (LY2090314) displaying disrupted morphology with cellular debris in H&E sections and marked upregulation of RHOC protein (intense brown staining) throughout the organoid epithelium. The dramatic RHOC induction correlates with activation of the non-canonical WNT/planar cell polarity pathway that executes apoptosis in hyperactivated WNT ("over-WNTed") tumor cells. Scale bars = 50 μm. (F,G) Rescue of GSK3i-induced growth inhibition by ROCK inhibition. (F) Representative brightfield images of AKP organoids treated with GSK3i (100 nM) ± ROCK1/2 inhibitor (Y-27632, 10 μM) for 72 hours. Scale bar: 100 μm. (G) Quantification of organoid growth by resazurin assay. n=4 biological replicates, mean ± SD. **p<0.01, one-way ANOVA. (H,I) Effect of ROCK inhibition on metastatic potential. (H) Representative gross brightfield (top) and tdTomato fluorescence (bottom) images of liver metastases 5 weeks after splenic injection. Left: AKPST organoids. Right: GSK3A/B knockout KPST organoids. Both organoid lines were cultured continuously with ROCK inhibitor (Y-27632, 10 μM) prior to injection. Scale bar: 5 mm. (I) Quantification of liver tumor burden calculated as Σ(mean fluorescence intensity × tumor area). n=5 mice per group, representative of 2 independent experiments. *p<0.05, unpaired t-test. (J) Genetic validation of RHOC-ROCK pathway. Growth curves of organoids with GSK3i dose response: AKPVT (control), AKPVT *Rhoc*^-/-^ (RHOC knockout), and AKPVT *Rock1*^-/-^/*Rock2*^-/-^ (ROCK1/2 double knockout). Growth measured by resazurin assay after 72 hours, normalized to untreated control for each genotype. n=3 biological replicates, mean ± SD. GSK3i = LY2090314 for all experiments. Data shown as mean ± SD. *p<0.05, **p<0.01.

**Figure S4:**
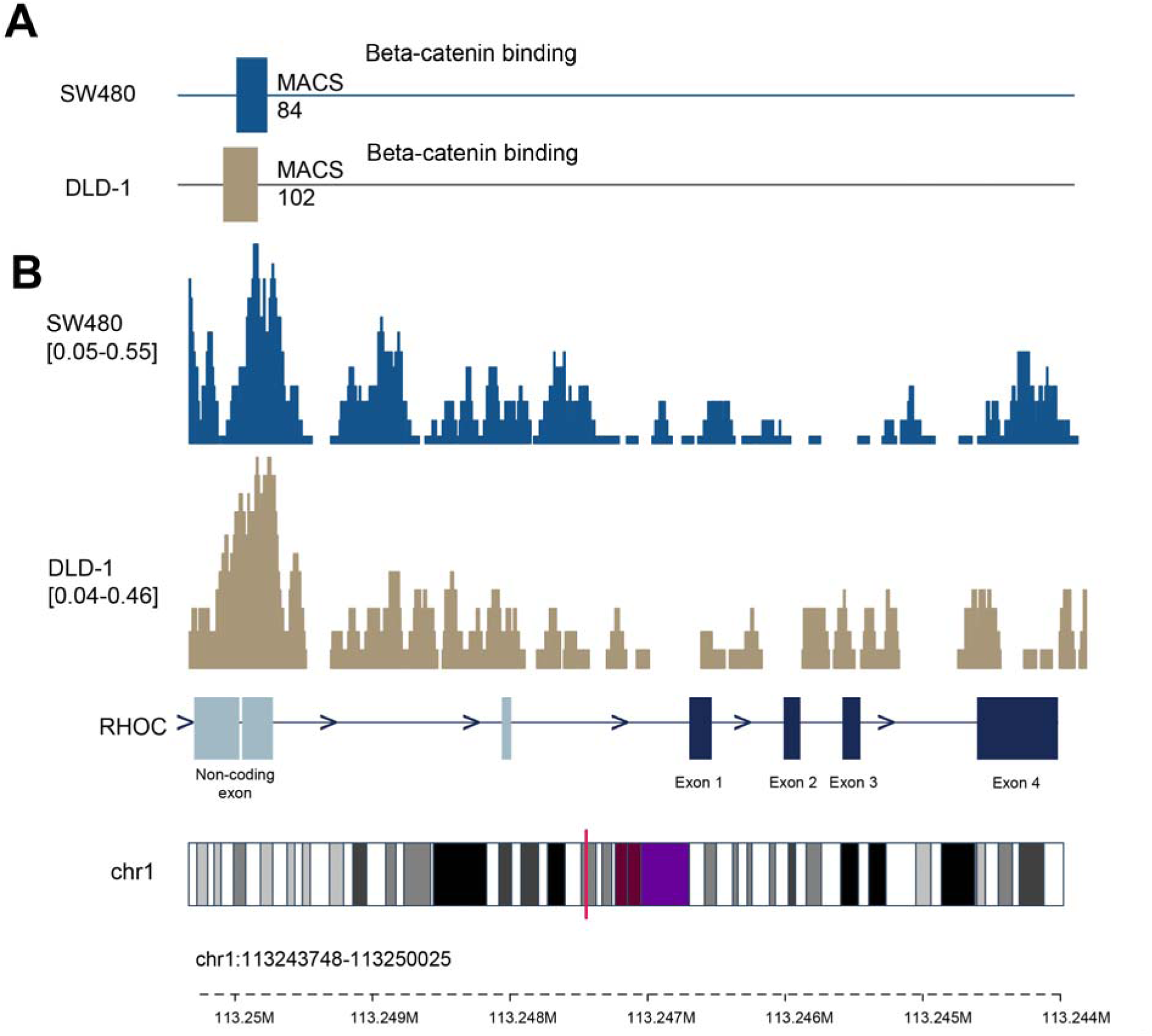
Beta-catenin binds directly to the *RHOC* locus in colorectal cancer cells. (A) ChIP-seq peak analysis showing beta-catenin binding at the *RHOC* locus in human colorectal cancer cell lines DLD-1 and SW480. Data obtained from publicly available datasets (DLD-1: SRX8929135; SW480: SRX8929141) and analyzed using ChIP-Atlas 3.0. Only peaks with MACS score >50 (q-value <1×10^-5^) are displayed. Peak heights represent normalized read density (reads per million). (B) Genome browser visualization of beta-catenin ChIP-seq signal at the *RHOC* locus. BigWig tracks show continuous signal intensity across the genomic region. Gene structure from UCSC genome browser (hg19 assembly) is shown below with genomic coordinates. Light blue boxes indicate untranslated regions (UTRs), dark blue boxes indicate coding exons, and lines represent introns. Arrows indicate transcription direction.

The identification of RHOC was unexpected. Through unbiased transcriptomic analysis, we identified this Rho family GTPase as one of the most significantly upregulated genes precisely at lethal WNT activation levels. To our knowledge, this is the first report linking RHOC activation to tumor cell death. As a Rho family GTPase, RHOC can be activated through the WNT/planar cell polarity pathway^57^, suggesting that extreme canonical WNT activation might "spill over" into non-canonical branches. To validate this at the protein level, we performed both immunofluorescence and immunohistochemistry analyses which confirmed dramatically elevated RHOC protein in GSK3 inhibitor-treated organoids. Immunohistochemical staining revealed minimal RHOC expression in untreated AKP organoids, while GSK3 inhibitor treatment induced striking RHOC upregulation throughout the organoid epithelium (Fig. 4D and E). The intense brown cytoplasmic staining pattern was particularly prominent in cells displaying disrupted morphology, mirroring the apoptotic changes observed with H&E staining. Both mouse and human CRC organoids showed this marked RHOC induction, with immunofluorescence analysis revealing increased cytoplasmic RHOC staining with prominent punctate accumulation, consistent with active GTPase localization patterns (Fig. 4D).

To establish the mechanistic link between WNT hyperactivation and RHOC induction, we analyzed published ChIP-seq datasets from colorectal cancer cells ^58^. This revealed direct beta-catenin binding peaks at the RHOC promoter region in both DLD-1 and SW480 CRC cell lines, with robust MACS scores (>50, q<1e-5) indicating high-confidence binding events (Fig. S4). The presence of multiple beta-catenin binding sites establishes RHOC as a direct transcriptional target of canonical WNT signaling—a connection further substantiated by our finding that GSK3 inhibitor treatment also increased beta-catenin expression itself. This beta-catenin-to-RHOC regulatory circuit creates a feed-forward amplification loop that ultimately triggers apoptosis.

We then investigated whether RHOC upregulation was functionally required for "over-WNTing"-induced cell death. RHOC signals through its downstream effector kinases ROCK1 and ROCK2, which are well-established mediators of apoptosis through cytoskeletal reorganization and nuclear disintegration.^59,60^ Remarkably, pharmacologic ROCK1/2 inhibition using GSK269962 almost completely rescued tumor organoids from GSK3 inhibitor-induced death without affecting baseline growth (Fig. 4F,G). The rescue was dose-dependent and specific, with maximal protection achieved at 100 nM ROCK inhibitor.

Having established the essential role of RHOC-ROCK signaling in over-WNTing-induced death, we next tested whether this mechanism operates in vivo. We leveraged the ROCK1/2 inhibitor rescue to generate triple knockout organoids (*Gsk3a*^-/-^, *Gsk3b*^-/-^, *Apc*^-/-^) in a KPST background. These organoids, which would normally die from constitutive "over-WNTing," could be maintained in culture only with continuous ROCK1/2 inhibition. Upon liver transplantation, these triple knockout organoids showed severely impaired engraftment and growth compared to control AKPST organoids, with >95% reduction in tumor burden (Fig. 4H,I). This demonstrates that the synthetic lethality between GSK3 loss and APC loss operates through the same RHOC-ROCK mechanism in vivo. To genetically establish the RHOC-ROCK axis as the death-executing mechanism, we generated CRISPR-mediated knockouts of RHOC and ROCK1/2 in AKP tumor organoids. Both RHOC knockout and ROCK1/2 double knockout organoids were completely resistant to "over-WNTing," maintaining robust growth even at GSK3 inhibitor concentrations that killed parental organoids (Fig. 4J). This genetic validation confirms that the RHOC-ROCK1/2 axis is not merely correlated with, but absolutely required for, WNT hyperactivation-induced tumor cell death.

Our mechanistic studies reveal an exquisite vulnerability in cancer cells: while they require high WNT signaling for survival, they exist perilously close to a threshold beyond which canonical WNT signaling spills into non-canonical pathways. This spillover activates RHOC-mediated planar cell polarity signaling, triggering ROCK1/2-dependent apoptosis—representing the first demonstration of RHOC as a bridge between canonical and non-canonical WNT signaling that can be exploited therapeutically. Normal cells, operating at lower baseline WNT levels, can tolerate substantial WNT enhancement without reaching this lethal threshold. This mechanistic understanding transforms "over-WNTing" from an empirical observation to a targetable synthetic lethal interaction, establishing a new paradigm where the driving oncogenic pathway becomes a fundamental vulnerability in cancer cells.

## DISCUSSION

Through functional organoid screening and genetic validation, we identified a specific GSK3 inhibitor that, as a sole agent, enhances intestinal regeneration by activating WNT signaling (Fig. 1). However, when applied to the precarious WNT-high state of tumor cells, the hyperactivation—or "over-WNTing"—tips them past a lethal threshold, eliminating neoplastic cells across the entire adenoma-carcinoma continuum (Fig. 2, S2). This differential response enabled marked therapeutic efficacy across all tested colorectal cancer models, from preventing adenoma formation to blocking metastatic seeding, achieving near-complete tumor elimination without toxicity (Fig. 3). The WNT hyperactivation state that was revealed engages a previously unknown signaling regime where canonical WNT pathway overstimulation spills into non-canonical planar cell polarity signaling, specifically inducing apoptosis in tumor cells (Fig. 4). Together, these findings establish a continuum model of WNT signaling that overturns fundamental assumptions in cancer therapy—demonstrating that hyperactivating oncogenic drivers can selectively kill tumor cells while simultaneously rejuvenating healthy tissue, representing the first anti-cancer treatment to achieve this dual benefit.

This unprecedented dual benefit arises from the irreversible nature of genetic APC loss, which creates a fundamental asymmetry between normal and transformed cells. While normal cells dynamically regulate WNT signaling and benefit from GSK3 inhibition, APC-mutant cells are genetically locked at a high setpoint which they cannot alter when pushed into lethal territory. This rigidity—essential for their niche independence—becomes their vulnerability, present in adenomas through metastatic cancer. Normal cells remain protected, operating safely below the danger threshold unless they undergo APC loss, which then provokes immediate elimination with concomitant WNT agonism. This self-regulating mechanism transforms the theoretical risk of general WNT enhancement into an asset—the treatment will not cause adenomas because it kills transformed cells the moment they arise, trough genetic APC loss, directly contradicting the initial assumption that WNT enhancement might increase cancer risk. Instead, our approach provides active protection against transformation while promoting tissue health. More broadly, our findings suggest that exploiting oncogenic inflexibility through pathway hyperactivation, rather than inhibition, may offer a generalizable strategy aligned to selectively exploit cancer’s fundamental biology The clinical implications of this intrinsic safety mechanism are profound, particularly for patients with familial adenomatous polyposis (FAP), who face colectomy in their twenties or thirties to avoid inevitable progression to CRC. Here, is a medical therapy capable of preventing adenoma progression while preserving and enhancing colonic function. The demonstration of survival extension in Apc^min^ mice and prevention of tumor initiation in temporally controlled models suggests that prophylactic administration could fundamentally alter the natural history of this disease, offering patients an alternative to organ loss. This approach is particularly compelling for the emerging epidemic of early-onset colorectal cancer, where patients under fifty face an especially cruel fate: surviving cancer only to endure decades of chemotherapy-related sequelae. Unlike conventional genotoxic agents that cause secondary malignancies, cardiovascular toxicity, and fertility compromise, "over-WNTing" achieves tumor elimination through a fundamentally different mechanism—pathway hyperactivation rather than DNA damage. For these younger patients facing decades of survivorship, a non-genotoxic therapy that preserves their long-term health represents a transformative advance in addressing this growing crisis.

Beyond these specific populations, the dual mechanism of tumor suppression with tissue regeneration enables therapeutic applications impossible with current approaches. In the adjuvant setting, where microscopic disease burden must be balanced against toxicity to normal tissues, the ability to administer therapy that simultaneously eliminates micrometastases while augmenting the intestinal stem cells and function could transform post-surgical care. Our demonstration that a single post-operative dose prevents metastatic seeding without observable toxicity suggests prophylactic use could become standard following primary resection. More broadly, the superior efficacy compared to 5-fluorouracil combined with absent toxicity compared to myelosuppression and mucositis, indicates potential both as monotherapy and in combination regimens where dose intensity limits use and outcomes.

The regenerative properties demonstrated in environmental enteropathy models reveal therapeutic potential beyond oncology. As an oral nanoparticle formulation, this approach could address the global burden of environmental enteropathy, where chronic intestinal damage perpetuates malnutrition and developmental stunting in millions of children worldwide. The ability to restore normal villus architecture and enhance nutrient absorption through epithelial regeneration represents a mechanistically novel intervention for a condition with no current pharmacological options. This same epithelial targeting mechanism positions WNT hyperactivation as the first potential therapy for inflammatory bowel disease to directly promote epithelial tissue healing rather than simply suppressing inflammation. Unlike current biologics that require systemic immunosuppression, local delivery of GSK3 inhibitor nanoparticles could enhance mucosal tissues while avoiding increased infection risk—particularly valuable in situations where tuberculosis and other opportunistic infections complicate immunosuppressive therapy.

The discovery that colon cancer cells exist at a precarious ’WNT-just-right’ state—not maximal activation—fundamentally reframes our understanding of oncogenic signaling. APC-mutant cells occupy a narrow signaling zone that maximizes proliferation and avoids apoptosis, but this precision creates an unexpected vulnerability. We demonstrated this constraint through multiple independent approaches— GSK3 inhibition, concentrated WNT proteins, and genetic manipulation—confirming this reflects fundamental tumor biology rather than compound-specific artifacts. By identifying LY2090314 as uniquely maintaining on-target activity at therapeutic concentrations, we validate GSK3 as a tractable therapeutic target when engaged with sufficient selectivity.

This work establishes hyperactivation of oncogenic drivers as a transformative therapeutic principle. In contrast to all current cancer therapies that compromise normal tissue function to achieve efficacy, WNT hyperactivation accomplishes selective tumor elimination while simultaneously promoting tissue regeneration. By elevating WNT signaling beyond the tolerance threshold of APC-mutant cells while enhancing normal stem cell function, we convert colorectal cancer’s essential driver mutation from an undruggable target into its primary vulnerability. This demonstrates how the constitutive signaling states that sustain cancer create inflexible dependencies that can be exploited therapeutically— establishing a new paradigm where cancer treatment enhances rather than compromises healthy tissue function.

### Limitations of the study

While our work establishes WNT hyperactivation as a therapeutic principle, important limitations remain. Our models capture key features of human colorectal cancer but lack the full immune microenvironment complexity, leaving interactions with immunotherapy unexplored. Although we identify RHOC-ROCK as the execution mechanism, the complete molecular network governing selective toxicity at high WNT levels remains to be elucidated. Safety studies span only four weeks; longer-term risks including crypt fission, ectopic WNT lesions, or extra-intestinal hyperplasia require investigation. The necessity of PLGA-encapsulation to achieve therapeutic window in vivo, combined with LY2090314’s rapid systemic clearance, presents pharmacokinetic challenges for clinical translation. Additionally, in vivo resistance mechanisms remain undefined. Understanding potential escape mechanisms will be essential for clinical development. Despite these limitations, the principle of exploiting oncogenic inflexibility through pathway hyperactivation opens new therapeutic avenues beyond colorectal cancer.

## RESOURCE AVAILABILITY

### Lead contact

Further information and requests for resources and reagents should be directed to and will be fulfilled by the lead contact, Omer H. Yilmaz.

### Materials availability

All unique reagents generated in this study are available from the corresponding author upon reasonable request.

### Data and code availability

#### Data and materials availability

RNA-seq data have been deposited at GEO and are publicly available as of the date of publication. Accession number: GSE273686 (token: Available to reviewers and final submission). All data reported in this paper are available in the main text or supplementary materials. This paper does not report original code. Any additional information required to reanalyze the data reported in this paper is available from the lead contact upon request.

## ACKNOWLEDGMENTS

We thank the Koch Institute Swanson Biotechnology Center for technical support, particularly the Hope Babette Tang (1983) Histology Facility, the Flow Cytometry Core, and the Preclinical Modeling Facility. We acknowledge the MIT BioMicro Center for sequencing support. This work was supported by National Cancer Institute grant K08CA277011 (G.E.), National Cancer Institute training grant T32CA009216 (G.E.), American Cancer Society grant CSCC-PF-23-996668-01-CSCC (G.E.), National Cancer Institute grant R01CA245314 (Ö.H.Y.), National Institute of Diabetes and Digestive and Kidney Diseases grant R01DK137971 (M.G.), and Department of Defense Peer Reviewed Cancer Research Program Idea Award W81XWH-19-1-0784 (M.G.). The content is solely the responsibility of the authors and does not necessarily represent the official views of the National Institutes of Health or the Department of Defense.

## AUTHOR CONTRIBUTIONS

G.E., J.B. and O.H.Y conceived the study. G.E. and J.B. designed and performed the experiments. G.E. and M.G. maintained the human colorectal cancer registry and enrolled patients for the study and analyzed the CHIP-Seq dataset. G.E. and J.B. wrote the paper. All the authors participated in the interpretation of the experiments, analyzed the data, and editing of the paper.

## DECLARATION OF GENERATIVE AI AND AI-ASSISTED TECHNOLOGIES

During the preparation of this work, the authors used Grammarly in order to proofread the text. After using this tool or service, the authors reviewed and edited the content as needed and takes full responsibility for the content of the publication.

## KEY RESOURCES TABLE

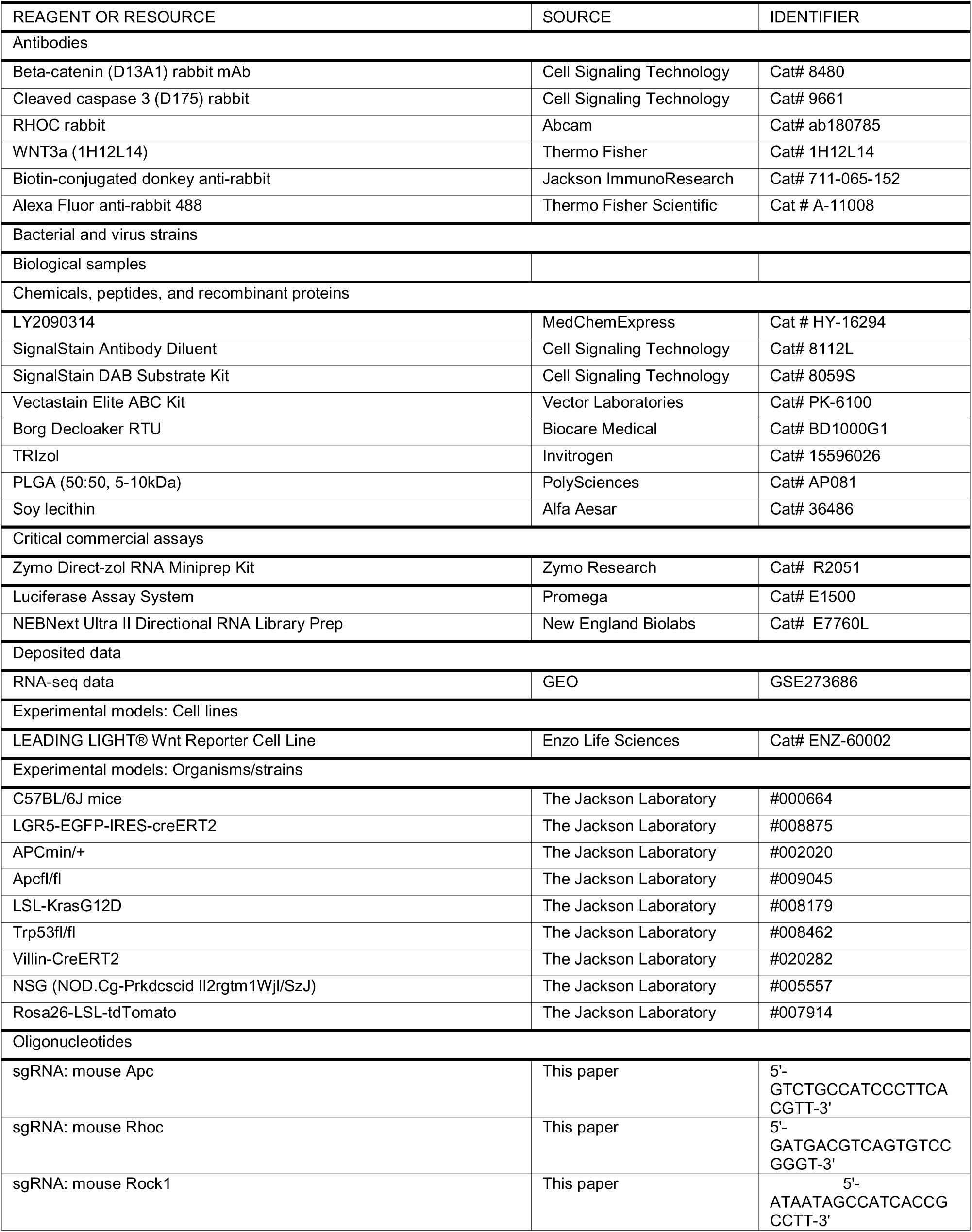
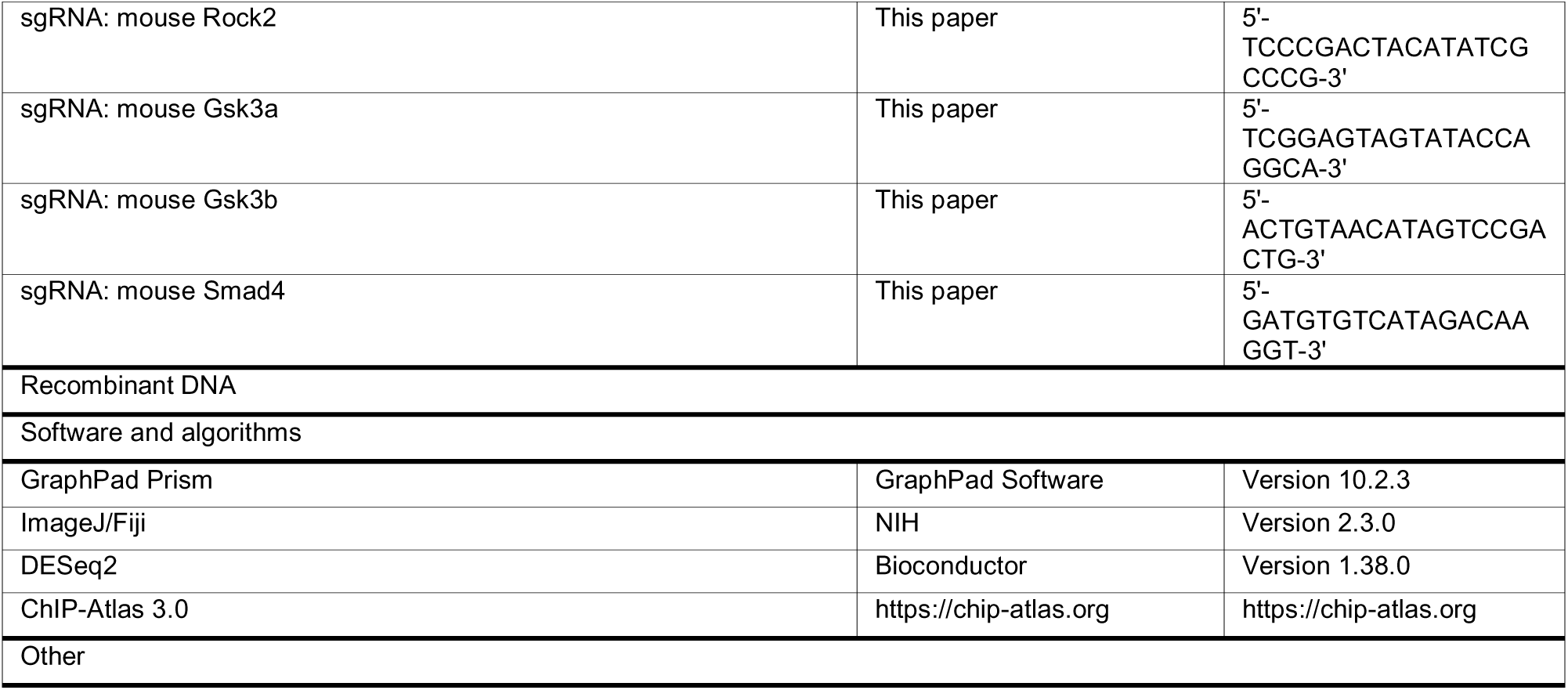

## EXPERIMENTAL MODEL AND STUDY PARTICIPANT DETAILS

### METHOD DETAILS

#### Materials and Methods

##### Ethics Statement

All mouse experiments were performed in accordance with protocols approved by the MIT Committee on Animal Care (Protocol #0123-001-26) and MGH Institutional Animal Care and Use Committee (Protocol #2019N000085). Human tissue collection was approved by the Mass General Brigham Institutional Review Board (Protocol #2018P002642). Written informed consent was obtained from all participants.

###### Mice

Mice were under the husbandry care of the Department of Comparative Medicine in the Koch Institute for Integrative Cancer Research. All procedures were conducted in accordance with the American Association for Accreditation of Laboratory Animal Care and approved by MIT’s Committee on Animal Care. Lgr5-2A-EGFP-2A-CreERT2, were used to assess stem cell functionality in response to GSK3 inhibitor treatment. Peritoneal carcinomatosis experiments used C57BL/6 organoids were performed on syngeneic C57BL/6 recipient mice. Orthotopic transplantation experiments with human CRC organoids utilized RAG2/IL2R mice. In this study, both male and female mice were used at the ages of 8-18 wks. Doxycycline was administered via food pellets (625 mg/kg) (Harlan Teklad, TD.09628). Nanoparticles (blank vs GSK3 inhibitor containing) were added to normal mouse diet (standard chow) of at a concentration of 50ug/4g (of LY2090314 GSK3 inhibitor) of chow and equivalent weight of non-LY2090314 containing blank nanoparticles. Mouse enemas of nanoparticles were performed using isofluorane anesthesia using a 20 gauge feeding needle (Fine Science Tools, 18064-20), where 20 ug of encapsulated LY2090314 was administered in each dose, 200 microliters, compared to blank PBS containing nanoparticles. For the Regional Basic Diet induced environmental enteropathy model, mice were weaned onto the formulated diet (Research Diets D09081701), which has significantly decreased protein, moderately decreased fat and minerals, compared to the isocaloric OpenStandard diet control, with ad libitum access to food.^43,46^ For induction of Cre-mediated recombination of *Apc* fl/fl; Villin CreERT2; *Rosa26* ^LSL-tdTomato^ mice, 100 μl of 1 mg/ml tamoxifen (MedChem Express, HY-13757A) in corn oil were injected intraperitoneally once. The following mouse strains were used, Lgr5-2A-EGFP-2A-CreERT2^48^, *Trp53* fl/fl,^61^ *Kras* ^LSL-G12D^,^62^ *Rosa26* ^LSL-ZsG^,^63^ *Rosa26* ^LSL-tdTomato63^, *Apc* fl/fl,^64^ Villin CreERT2^65^, APC min^55^, *Rag2 Il2r* (Taconic) were all maintained on a C57BL/6J background.

###### Organoid generation and culture

Colon organoids were generated by blunt dissection of mouse colons (8-16 weeks old). Fecal material was removed and the colon tissues were rinsed in ice cold PBS, then cut into three 1.5 cm length pieces. They were incubated in PBS with 5 mM EDTA at 37°c for 15 min, with constant agitation. The liberated crypts were then centrifuged at 1000 RCF for 10 s, resuspended in media and Matrigel (Corning, 356231) and plated onto tissue culture plastic dishes, in 10 microliter volumes, permitted to gel at 37°c, then overlaid with culture media and cultured in a tissue culture incubator (5% CO2, 100% humidity at 37°c). Normal mouse organoids were maintained in Advanced DMEM/F12 (Gibco,12634010), supplemented with 5% FBS (Hyclone, SH30071), penicillin/streptomycin (Gibco, 15140122), and 2mM GlutaMAX (Gibco, 35050061), EGF 50 ng/ml (Peprotech, 315-09) and 50% WRN supernatant.^31,32^ Colon cancer organoids were typically maintained in Advanced DMEM/F12 (Gibco,12634010), supplemented with 5% FBS (Hyclone, SH30071), penicillin/streptomycin (Gibco, 15140122), and 2mM GlutaMAX (Gibco, 35050061). Human organoids of normal tissue, adenoma, and cancers were generated under MassGeneralBrigham IRB protocol (2013P002559). Written informed consent was obtained from all participants. Human cancer organoids were embedded in Matrigel and cultured in media (Advanced DMEM supplemented with 5% FBS (Hyclone, SH30071), 50 ng/ml EGF (PeproTech, 315-09), 500 nM A83-01 (Tocris, 2939), 10μM SB202190 (Sigma-Aldrich, S7067), PGE2 500nM (Cayman Chemical 14010), 10 μM nicotinamide (Sigma-Aldrich, N3376), 1 μM N-acetyl-l-cysteine (Sigma-Aldrich, A7250), B27 (Thermo Fisher Scientific, 17504044), penicillin/streptomycin (Gibco 15140122), and GlutaMAX (Gibco, 35050061). Human normal organoids were maintained in the same media as human cancer organoids but also with 50% WRN. In general, organoid cultures were passaged weekly, with 1:6 split ratio, where matrigel droplets were mechanically dissociated and collected, centrifuged at 1000 RCF for 1 minute, treated with TrypLE (Gibco12604013) for 1 min at 37c with constant agitation, re-centrifuged, TrypLE removed, and cells were resuspended in media and Matrigel, plated on tissue culture plastic, using 10 microliter droplets, similar to the initial primary crypt culture.

###### Patient-derived organoid culture

Human colorectal tissue was obtained from patients undergoing colonoscopy or surgical resection at Massachusetts General Hospital under IRB protocol #2018P002642. Normal tissue was obtained from macroscopically normal regions >10 cm from any tumor. Tumor tissue was confirmed by pathological examination. Tissue was processed within 2 hours of collection.

###### Organoid growth assays

Organoid viability and growth were quantified using resazurin metabolic assay (Alamar Blue Active Ingredient), a non-destructive method that measures viable cell number without requiring cell lysis. This assay exploits the fundamental principle that metabolically active cells maintain a reduced intracellular environment through the electron transport chain and cytoplasmic reductases. Resazurin (7-Hydroxy-3H-phenoxazin-3-one 10-oxide), a blue non-fluorescent redox indicator with an oxidation-reduction potential of +380 mV, readily penetrates cell membranes and is reduced by viable cells to resorufin (7-Hydroxy-3H-phenoxazin-3-one), which exhibits bright pink fluorescence (λEx/λEm: 530-560/590 nm).

The reduction occurs through multiple cellular mechanisms including mitochondrial, cytoplasmic, and microsomal dehydrogenases/reductases, with the rate of reduction directly proportional to the number of metabolically active cells. Unlike tetrazolium-based assays (e.g., MTT), resazurin reduction produces a soluble fluorescent product that diffuses into the culture medium, eliminating the need for cell lysis and organic solvent extraction. This non-invasive approach enables kinetic monitoring and preserves organoid architecture for downstream analyses.

Stock solutions of Resazurin (Sigma-Aldrich, R7017) were prepared at 0.5mg/ml in PBS and sterile filtered. At experimental endpoints, resazurin was added to culture wells at a final concentration of 50 μg/ml (0.2 mM) and incubated for 2 hours at 37°C, 5% CO2. During incubation, the continuous metabolic activity of viable cells drives the reduction of resazurin, while non-viable cells lacking functional metabolism cannot catalyze this conversion. Fluorescence measurements were acquired using a microplate reader (excitation 560 nm/emission 590 nm). Background fluorescence was corrected by subtracting measurements taken immediately upon resazurin addition, accounting for any intrinsic fluorescence.

The fluorescence intensity demonstrates linear correlation with viable cell number across a dynamic range of 10²-10L cells per well, as established by calibration curves. For organoid growth assays, dissociated organoid cells were seeded at 1000 cells/10μL Matrigel droplet and cultured for 4 days prior to viability quantification. Data are presented as relative fluorescence units (RFU) normalized to control conditions, providing a quantitative measure of relative viable cell numbers. The high sensitivity (detection limit ∼100 cells) and broad linear range make this assay ideal for monitoring organoid growth dynamics across different experimental conditions.

###### GSK3 inhibitor small molecule screen

The following GSK3 inhibitors were obtained and prepared according to manufacturer’s specifications: LY2090314 (Cayman Chemical, 22211), CHIR99021 (LC Laboratories, C-556), A1070722 (Sigma-Aldrich, SML0863), QS11 (Cayman Chemical, 21247) 9-ING-41 ((MedChem Express, HY-113914), SB216763 (Cayman Chemical, 10010246) BIM I (Cayman Chemical 13298), BIM IX (Cayman Chemical,13334), Tideglusib (Cayman Chemical, 16727), BIO (Cayman Chemical, 13123) CHIR98014 (Cayman Chemical, 15578) and in general were dissolved in DMSO. These compounds were tested on normal colon and GSK3 alpha and beta knockout organoids across a dose response in the absence of any other WNT agonists and growth was measured by resazurin assay.

###### Organoid editing

To generate GSK3A/B double knockout organoids, wildtype normal colon organoids were transfected with a hCas9 dual guide targeting plasmid, sg*GSK3A* (5′-TCGGAGTAGTATACCAGGCA-3′) and sg*GSK3B* (5′-ACTGTAACATAGTCCGACTG-3′), and double knockouts were selected by growth upon WNT agonist withdrawal. To generate *Rock1/2* double knockout organoids, AKP colon cancer organoids were transfected with a hyPBase expression plasmid at a 1:1 molar ratio with a PiggyBac plasmid expressing dual guide, sg*Rock1* (5′-ATAATAGCCATCACCGCCTT-3’), and sg*Rock2* (5′-TCCCGACTACATATCGCCCG-3′), hCas9, and PuroR. *Rock1/2* double knockouts were selected using puromycin at 4 ug/ml. To generate RHOC knockout organoids, AKP colon cancer organoids were transfected with a hyPBase expression plasmid at a 1:1 molar ratio with a PiggyBac plasmid expressing guide, sg*Rhoc* (5′-GATGACGTCAGTGTCCGGGT-3’), hCas9, and BsdR.

To generate *Apc*^-/-^ tdTomato organoids, colon organoids from *Rosa26* ^LSL-tdTomato^ mice were transfected with pSECC plasmid ^66^ containing Cre, Cas9, and sg*Apc* (5’-GTCTGCCATCCCTTCACGTT-3′), and organoids were selected by WNT agonist withdrawal. To generate AKP and AKP tdTomato organoids, colon organoids from *Kras*^LSL-G12D^ +/-; *Trp53* ^fl/fl^ and *Kras*^LSL-G12D^ +/-; *Trp53*^fl/fl^, *Rosa26*^LSL-tdTomato^ were transfected with pSECC plasmid containing Cre, Cas9, and sg*Apc* (5’-GTCTGCCATCCCTTCACGTT-3′), and organoids were selected by WNT agonist withdrawal, addition of 10 micromolar Nutlin-3 (Sigma-Aldrich, N6287), and 1 micromolar Gefitinib (Cayman Chemical, 13166). To generate AKPS tdTomato organoids, colon organoids from *Apc*^fl/fl^; *Kras*^LSL-G12D^ +/-; *Trp53*^fl/fl^, *Rosa26*^LSL-tdTomato^ mice were transfected with pSECC containing Cre, Cas9, and sg*Smad4* (5’-GATGTGTCATAGACAAGGT -3’), and organoids were selected by WNT agonist withdrawal, addition of 10 micromolar Nutlin-3, 1 micromolar Gefitinib (Cayman Chemical, 13166) and TGFB 10ng/ml (Peprotech, 100-21C). To generate GSK3A/B AKPST organoids, colon organoids from *Apc^f^*^l/fl^; *Kras*^LSL-G12D^ +/-; *Trp53*^fl/fl^, *Rosa26*^LSL-tdTomato^ mice were transfected with a hCas9 dual guide targeting plasmid, sg*Gsk3a* (5′-TCGGAGTAGTATACCAGGCA-3′) and sg*Gsk3b* (5′-ACTGTAACATAGTCCGACTG-3′), and were selected with WNT agonist withdrawal. These resultant GSK3A/B double knockout, *Apc*^fl/fl^; *Kras*^LSL-G12D^ +/-; *Trp53*^fl/fl^, *Rosa26*^LSL-tdTomato^ organoids were then grown with the addition of 100nM GSK269962 ROCK1/2 inhibitor (Cayman Chemical, 19180) and 500nM A83-01 (Cayman Chemical, 9001799) and transfected with pSECC-sgSMAD4, and were selected with addition of 10 micromolar Nutlin-3, 1 micromolar Gefitinib (Cayman Chemical, 13166) and TGFB 10ng/ml (Peprotech, 100-21C). APC excision was confirmed by PCR genotyping for the excised allele, (forward 5’-GTAGTGCACACCTTTAATCCCTTCTC - 3’, reverse 5’ - TACGCTTCTAGAGCAGCAAACTTAC - 3’), and absence of the pre-excision allele, (forward 5’ - CTAGTACTTTTCAGACGTCATG - 3’, reverse 5’ - CCCACCTCCACTTTCAATAA - 3’). AKP tdTomato organoids were alternatively generated from colon organoids from *Apc*^fl/fl^; *Kras*^LSL-G12D^ +/-; *Trp53*^fl/fl^, *Rosa26*^LSL-tdTomato^ Villin-CreERT2 mice, treated with 4-hydroxytamoxifen at 100ng/ml for 3 days, and selected WNT agonist withdrawal, addition of 10 micromolar Nutlin-3, and 1 micromolar Gefitinib. ZsGreen^+^ organoids with no cancer mutations were generated from organoids from Rosa26^LSL-ZsGreen^ mice transduced with Ad5CMVCre (UI Viral Vector Core, Iowa-5). To generate WNT3a expressing organoids, mice harboring the following genetic germline variants were maintained: *Kras*^G12D^, *Trp53* ^fl/fl^, Villin-^CreERT2^, *Rosa26*-^LSL-tdTomato^. These organoids were isolated from colon crypts and exposed to 4-hydroxy-tamoxifen at 1uM for 3 days and maintained in Gefitnib (10 uM) and Nutlin-3(10 μM). The resulting KPT mutant organoids were then transfected with WNT3a transgene using a piggybac transposon mediated integration where helper plasmid was provided to the transgene using 15 µl of Lipofectamine 2000 reagent in 250 microliters of Opti-MEM medium (Thermo Fisher Scientific, 31985062), 3 ug of transgene and 3 ug of PiggyBac helper plasmid high activity HyPBase (Vectorbuilder). The two vectors were mixed and added to a single cell suspension of the organoid cells and spun at 100g for 1 hour. Cells were selected for puromycin resistance using puromycin at 4ug/ml for one week, after replating in Matrigel and waiting 1 day, and individual clones were picked and maintained for experiments. The TOP/tdTomato reporter was designed similarly to the Super 8x TOPFlash reporter (Addgene #12456), with TCF/LEF DNA binding sites sequence: AGATCAAAGG, (total of 13 TCF/LEF Binding sites) placed upstream a minimal P promoter and the tdTomato gene. EF1A was used after the tdTomato to drive NeoR. PiggyBac ITR sequences flanked these elements. This was transfected into wildtype normal colon mouse organoids, AKP colon cancer mouse organoids and human CRC organoids, by mixing the TOP/tdTomato plasmid (3ug) with (3ug) of the PiggyBac helper plasmid in Lipofectamine 2000, all in 250 microliters of Opti-MEM. The organoid cells were first dissociated to single cells by mechanical trituration and then 3 minutes of exposure to TrypLE. After addition of the TOP/tdTomato plasmid and the PiggyBac helper plasmid in Lipofectamine 2000 to the cells, the mixture was centrifuged at 100g for 1 hour, and the resulting cells were then embedded in Matrigel and grown. G418 was used at 1mg/ml for 2 weeks to select for plasmid containing organoids. sh*Apc* plasmid was transduced into organoids using standard lentiviral production and transduction. The plasmid was a generous gift from Lukas Dow.^67^ For lentivirus production, 10 µl of Lipofectamine 2000 reagent in 250 µl Opti-MEM medium (Thermo Fisher Scientific, 31985062) and a total of 4 µg of DNA (2 µg of lentiviral backbone constructs, 1 µg of pSpax2 (addgene, #12260), and 1 µg of pMD2.G (addgene, #12259)) in 250 µl Opti-MEM medium were mixed together, incubated for 15 min, and added to 293T cells cultured in 6-well plates. Culture media were changed next day. The supernatant was collected 48 hours after the media change and passed through a 0.45-μm filter and immediately used. For lentivirus transduction, organoids were dissociated by incubation with TrypLE Express for 5 min at 37°C, transduced with the lentivirus supernatant by spinoculation at 600g at 32 °C for 60 min in a 48-well culture plate, and incubated for another 4 hr in an incubator at 37°C. Organoids were then embedded in Matrigel and cultured in Advanced DMEM/F12 supplemented with 5% FBS penicillin/streptomycin and 2mM GlutaMAX. Two days after transduction, 4 μg/ml puromycin was added to the culture media to select for organoids containing the plasmid.

###### WRN concentration procedure

To generate concentrated WNT proteins, supernatant from a Wnt3a, R-Spondin3, and Noggin (WRN) secreting cell line was used.^31^ Wnt proteins require palmitoylation for maintenance of functional activity, and resuspension of recombinant expressed proteins often require the use of detergents such as CHAPS, which are highly toxic when used at high concentration.^50^ To avoid this we concentrated WRN supernatant using PEG-8000 mw (Sigma Aldrich P2139). Briefly, 160g of PEG-8000, 28g of NaCl and 40ml of 10x PBS was dissolved in 18 Mohm water, and pH was adjusted to 7.4. The solution was filtered using 0.45 micron filter (Millipore S2HVU05RE). After 24 hours of production from a confluent 150mm cm^2^ tissue culture treated dish (Genesee 25-203) 37.5 ml of supernatant from the L-WRN cell line was collected and centrifuged at 1000 RCF for 10 minutes, and filtered through a 0.2 micron filter (Millipore S2GVU05RE). 12.5 ml of the PEG solution was added to 37.5 ml of supernatant, shaken for 60 sec and placed on nutating rotation overnight at 4°C. The solution was then centrifuged at 3000 RCF for 1 hr at 4°C. The supernatant was removed, and the pellet was resuspended in 1.9ml of culture media to generate a 20x concentrated solution.

###### Organoid implantation models

Peritoneal injections. Mouse organoids were implanted into syngeneic mice for the peritoneal carcinomatosis assay as follows: Mouse organoid cells harboring mutations for *Apc*, *Kras*, and *Tp53*, as well as constitutive expression of tdTomato, were dissociated from day 4 growth Matrigel droplets. The organoids were separated from their Matrigel domes by centrifugation at 1000g for 60 sec, and exposed to TrypLE for 3 min and passed through a 70 micron filter to obtain a single cell suspension in PBS. 150,000 dissociated organoids cells were injected into the peritoneums of mice, in a total volume of 500 microliters per injection, in PBS. The cells were permitted to implant for 3 days and treatments as specified were initiated. For human carcinoma organoids, human tumor organoids were permitted to implant for 7 days prior to initiation of therapy. At experimental endpoint, tumors were dissected from the peritoneal cavity and imaged. For injections with beta-cyclodextrin (Sigma Aldrich, C4767), 20% w/w beta-cyclodextrin was used as a carrier in which LY2090314 was dissolved at 0.1mg/ml, with 500 microliters injected per dose. Nanoparticle GSK3 was dosed at 500 microliter volume, containing 50 micrograms of encapsulated LY2090314. Images of tumors were acquired on a Nikon SMZ18 stereoscopic dissection microscope with a DS-Fi3 camera, and a Lumencor light engine as a fluorescent light source. Tumors were analyzed using ImageJ 1.54f.

###### Splenic injections

For splenic injections, mice were anesthetized using isofluorane and maintained under surgical sterile conditions. After surgical skin preparation, a small incision in the left lateral flank was made using iris scissors, and 200ul of tumor cells was infiltrated into the spleens of the mice over a 5 min period. The cells were prepared similarly to the peritoneal injection model with the associated cell concentrations modified as listed. After the infusion, the spleens were carefully removed using thermal cautery, and the wound was closed using 5-0 absorbable sutures on the inner layer, followed by 5-0 polypropylene sutures on the exterior layer, and removed after 1 week post-operatively. When challenged with immediate chasing of nanoparticles, a dose of 10 ug (LY2090314) of GSK3 inhibitor encapsulated nanoparticles (100 microliter solution) was injected immediately after infusing tumor cells after a 5 minute wait period, compared to blank nanoparticle controls. At experimental endpoint, tumors were dissected from the peritoneal cavity and imaged. Images of tumors were acquired on a Nikon SMZ18 stereoscopic dissection microscope with a DS-Fi3 camera, and a Lumencor light engine as a fluorescent light source. Tumors were analyzed using ImageJ 1.54f.

###### TOP Flash WNT Assay

The TOP/Flash assay was performed as per protocol from the Promega luciferase assay system kit (E6110). The cell line used was the Enzo Leading light WNT reporter cell line, ENZ-61001. Briefly, the cells were maintained in standard advanced DMEM/F12 media with antibiotics, Glutamax and 5% FBS, passaged twice weekly. The cells were plated 2 days prior to assay initiation and grown to near confluence. WNT media (either recombinant proteins or GSK3 inhibitors) were then applied to the cells in the media, and luciferase activity was assessed at 24 hours of exposure using a Tecan spectrophotometer, set for luminescence detection (TECAN infinite 200Pro).

###### Immunohistochemistry (IHC) and immunofluorescence (IF)

Tissues were fixed in 10% formalin, paraffin embedded and sectioned in 4-5 micron sections as previously described.^68^ Antigen retrieval was performed using Borg Decloaker RTU solution (Biocare Medical, BD1000G1) with a pressurized Decloaking Chamber (Biocare Medical, NxGen). All antibody dilutions were performed in SignalStain Antibody Diluent (CST, 8112L). For immunofluorescence, the following primary antibodies were used: WNT3a (1H12L14) 1:500 (Thermo Fisher) and RHOC 1:500 (Abcam, ab180785). Alexa Fluor secondary antibodies, anti-goat 488, anti-rabbit 488, anti-goat 568, anti-rabbit 568 (1:500, Thermo Fisher Scientific), were used for visualization. Slides were stained with DAPI for 10 min and covered with MOWIOL with DABCO mounting media. Images were acquired using a Nikon Eclipse 90i upright microscope equipped with a Hamamatsu Orca-ER CCD camera, and APC line 1200 light source. Whole organoid images, both brightfield and fluorescence, were acquired on a Keyence BZ-X series microscope, Z-series, and applying the full focus function to generate flattened images. For immunohistochemistry, the following primary antibodies were used: beta-catenin (D13A1, Cell Signaling Technology #8480, 1:500), cleaved caspase 3 (D175, Cell Signaling Technology #9661, 1:500), and RHOC (Abcam #ab180785, 1:500). Biotin-conjugated donkey anti-rabbit secondary antibody (Jackson ImmunoResearch #711-065-152, 1:500) was applied, followed by Vectastain Elite ABC immunoperoxidase detection kit (Vector Laboratories #PK-6100). Visualization was performed using SignalStain DAB Substrate Kit (Cell Signaling Technology #8059S).

###### Image Analysis and Quantification

Immunohistochemistry quantification was performed using Fiji/ImageJ. For cleaved caspase 3 analysis, images were acquired at 10x magnification and processed using color deconvolution to separate hematoxylin and DAB channels. Nuclei were segmented from the hematoxylin channel using Otsu thresholding followed by watershed separation. CC3 intensity was measured as 255 minus mean DAB intensity per cell, with cells classified as CC3-positive if mean DAB intensity was below 120. A total of 3,797 control cells (13 fields, 3 biological replicates) and 4,677 treated cells (14 fields, 3 biological replicates) were analyzed.

For nuclear beta-catenin quantification, images were acquired at 20x magnification and processed using color deconvolution to separate hematoxylin and DAB channels. Both channels were median filtered (radius = 3 pixels). Nuclear segmentation was performed on the hematoxylin channel using Bernsen’s adaptive local thresholding (radius = 25 pixels), followed by hole filling and watershed separation. Nuclear beta-catenin intensity was measured as mean DAB intensity within nuclei ≥1500 pixels². A total of 805 control cells (10 fields, 3 biological replicates) and 1,608 treated cells (13 fields, 3 biological replicates) were analyzed. Statistical analysis was performed using unpaired t-tests in GraphPad Prism 10.2.3. Detailed analysis macros are available upon request.

###### RNA extraction, RNA-seq library preparation, processing and differential expression analysis

RNA was extracted from organoids using TRIzol and Zymo Direct-zol RNA Miniprep Kits (Zymo Research, CA, USA) according to the manufacturer’s instructions. For RNA-seq library preparation, NEBNext Ultra II Directional RNA preparation with poly(A) selection is used to selectively capture mRNA from total RNA and barcoded libraries were generated. 2 x 40 paired-end sequencing was performed on an Illumina NextSeq 500. RNA-seq data processing and differential expression analysis was performed by the Koch Institute sequencing core.

RNA-seq data from GEO (Number to be assigned, metadata file and data were uploaded 7/29/2024) were used to quantify transcripts from the mm10 mouse assembly with the Ensembl version 101 annotation using Salmon version 1.3.0^69^and gene level summaries were prepared using tximport version 1.16.1.^70^. Differential expression analysis was done with DESeq2 version 1.30.1^56^ and differentially expressed genes were defined as those having an absolute log2 fold change greater than 1 and an adjusted P value less than 0.01.^71^

###### Nanoparticle Formulation and Synthesis

Poly(lactic-co-glycolic acid) (PLGA) nanoparticles loaded with LY2090314 were generated via a double emulsion technique.^68^ Briefly, 0.5 g of PLGA (LG 50:50 Mn 5,000-10,000 Da acid endcap, Polyscitech AP081) was dissolved in 2.5 ml of dichloromethane, and a LY2090314 solution (10mg/ml) 0.5 ml was added to the solution and sonicated for 30 sec (Hanchen Instrument 300w 3mm probe). The emulsion was then transferred to 50 ml of cold PBS with soy lethicin (Alfa Aesar 36486) (10 mg / ml) and was sonicated for 30s twice, with a 30 sec wait interval. The dichloromethane was then removed from the solution via stirring for 12 hrs at 4 c. The particles were isolated and washed via centrifugation (5,000 g for 30 m x 3 times) and frozen for long term storage.

###### Nanoparticle characterization

Dynamic light scattering measurements were performed using a Wyatt DynaProII Plate Reader at 25°C. Particles were diluted in 18 MΩ water to a concentration of 0.1 mg/mL. Size distribution was analyzed using DYNAMICS software version 7.8.

###### Analysis of Beta-catenin ChIP-seq data from CRC cell lines

Publicly available Beta-catenin ChIP-seq data were obtained for DLD-1 (SRX8929135) and SW480 (SRX8929141) cell lines from prior publication and analyzed on the ChIP-Atlas 3.0 platform (PMID: 38749504).^58^ Briefly, histone-DNA complexes were isolated with 19505943 -beta-catenin antibody (Cell Signalling, #9581). Illumina sequencing libraries were prepared from the ChIP and Input DNAs by the standard consecutive enzymatic steps of end-polishing, dA-addition, and adaptor ligation. After a final PCR amplification step, the resulting DNA libraries were quantified and sequenced on the NextSeq 500 platform (Illumina) with 100 bp single-end reads. Fastq files were aligned to the hg19 reference genome with Bowtie 2.^72^ The generated SAM files were converted to BAM files with samtools v0.1.19^73^ with duplicate reads removed. Bedtools v 2.17^74^ was used to generate BedGraph-formatted scores in reads per million mapped units. BedGraph files were subsequently binarized into BigWig format, and peak calls with q < 1e-5 were made with MACS2.^75^

### QUANTIFICATION AND STATISTICAL ANALYSIS

Unless otherwise specified in the figure legends or Method Details, all experiments reported in this study were repeated at least three independent times. All sample number (n) of biological replicates and technical replicates, definition of center, and dispersion and precision measures can be found in the figure legends. The images for H&E, immunofluorescence, and immunohistochemistry represent one of 3 biological replicates unless otherwise stated. All values are presented as mean ± SD unless otherwise stated. Intergroup comparisons were performed using unpaired two-tailed t-tests or one-way analysis of variance (ANOVA) with post-hoc Tukey’s multiple comparison. P values of < 0.05 were considered to be significant. Statistical analysis was performed by GraphPad Prism 10.2.3. No samples or animals were excluded from analysis. Age- and sex-matched mice were randomly assigned to groups. Studies were not conducted blind. Please note that statistical details are found in the figure legends. Sample sizes were determined based on pilot experiments and previous publications. For organoid experiments, n=3-4 biological replicates provided >80% power to detect a 50% difference in growth with α=0.05. For in vivo experiments, n=5-6 mice per group provided >80% power to detect a 40% reduction in tumor burden with α=0.05.

## REFERENCES

1. Bienz, M., and Clevers, H. (2000). Linking Colorectal Cancer to Wnt Signaling. Cell 103, 311–320. 10.1016/S0092-8674(00)00122-7.

2. Fearon, E.R., and Vogelstein, B. (1990). A genetic model for colorectal tumorigenesis. Cell 61, 759– 767. 10.1016/0092-8674(90)90186-I.

3. Fevr, T., Robine, S., Louvard, D., and Huelsken, J. (2007). Wnt/β-Catenin Is Essential for Intestinal Homeostasis and Maintenance of Intestinal Stem Cells. Molecular and Cellular Biology 27, 7551– 7559. 10.1128/MCB.01034-07.

4. Krausova, M., and Korinek, V. (2014). Wnt signaling in adult intestinal stem cells and cancer. Cellular Signalling 26, 570–579. 10.1016/j.cellsig.2013.11.032.

5. Fodde, R. (2002). The APC gene in colorectal cancer. European Journal of Cancer 38, 867–871. 10.1016/S0959-8049(02)00040-0.

6. Takahashi-Yanaga, F., and Kahn, M. (2010). Targeting Wnt Signaling: Can We Safely Eradicate Cancer Stem Cells? Clinical Cancer Research 16, 3153–3162. 10.1158/1078-0432.CCR-09-2943.

7. Zhao, H., Ming, T., Tang, S., Ren, S., Yang, H., Liu, M., Tao, Q., and Xu, H. (2022). Wnt signaling in colorectal cancer: pathogenic role and therapeutic target. Mol Cancer 21, 144. 10.1186/s12943-022-01616-7.

8. Schatoff, E.M., Leach, B.I., and Dow, L.E. (2017). WNT Signaling and Colorectal Cancer. Curr Colorectal Cancer Rep 13, 101–110. 10.1007/s11888-017-0354-9.

9. Bahrami, A., Amerizadeh, F., ShahidSales, S., Khazaei, M., Ghayour-Mobarhan, M., Sadeghnia, H.R., Maftouh, M., Hassanian, S.M., and Avan, A. (2017). Therapeutic Potential of Targeting Wnt/β-Catenin Pathway in Treatment of Colorectal Cancer: Rational and Progress: Wnt P ATHWAY IN CRC. J. Cell. Biochem. 118, 1979–1983. 10.1002/jcb.25903.

10. Lyou, Y., Habowski, A.N., Chen, G.T., and Waterman, M.L. (2017). Inhibition of nuclear Wnt signalling: challenges of an elusive target for cancer therapy. British J Pharmacology 174, 4589– 4599. 10.1111/bph.13963.

11. Chen, G.T., Tifrea, D.F., Murad, R., Habowski, A.N., Lyou, Y., Duong, M.R., Hosohama, L., Mortazavi, A., Edwards, R.A., and Waterman, M.L. (2022). Disruption of β-Catenin–Dependent Wnt Signaling in Colon Cancer Cells Remodels the Microenvironment to Promote Tumor Invasion. Molecular Cancer Research 20, 468–484. 10.1158/1541-7786.MCR-21-0349.

12. Jung, Y.-S., and Park, J.-I. (2020). Wnt signaling in cancer: therapeutic targeting of Wnt signaling beyond β-catenin and the destruction complex. Exp Mol Med 52, 183–191. 10.1038/s12276-020-0380-6.

13. Chang, L., Jung, N.Y., Atari, A., Rodriguez, D.J., Kesar, D., Song, T.-Y., Rees, M.G., Ronan, M., Li, R., Ruiz, P., et al. (2023). Systematic profiling of conditional pathway activation identifies context-dependent synthetic lethalities. Nat Genet 55, 1709–1720. 10.1038/s41588-023-01515-7.

14. Albuquerque, C. (2002). The “just-right” signaling model: APC somatic mutations are selected based on a specific level of activation of the beta-catenin signaling cascade. Human Molecular Genetics 11, 1549–1560. 10.1093/hmg/11.13.1549.

15. Langlands, A.J., Carroll, T.D., Chen, Y., and Näthke, I. (2018). Chir99021 and Valproic acid reduce the proliferative advantage of Apc mutant cells. Cell Death Dis 9, 255. 10.1038/s41419-017-0199-9.

16. Flanagan, D.J., Vincan, E., and Phesse, T.J. (2019). Wnt Signaling in Cancer: Not a Binary ON:OFF Switch. Cancer Research 79, 5901–5906. 10.1158/0008-5472.CAN-19-1362.

17. Christie, M., Jorissen, R.N., Mouradov, D., Sakthianandeswaren, A., Li, S., Day, F., Tsui, C., Lipton, L., Desai, J., Jones, I.T., et al. (2013). Different APC genotypes in proximal and distal sporadic colorectal cancers suggest distinct WNT/β-catenin signalling thresholds for tumourigenesis. Oncogene 32, 4675–4682. 10.1038/onc.2012.486.

18. Huels, D.J., Ridgway, R.A., Radulescu, S., Leushacke, M., Campbell, A.D., Biswas, S., Leedham, S., Serra, S., Chetty, R., Moreaux, G., et al. (2015). E-cadherin can limit the transforming properties of activating β-catenin mutations. EMBO J 34, 2321–2333. 10.15252/embj.201591739.

19. Gay, D.M., Ridgway, R.A., Müller, M., Hodder, M.C., Hedley, A., Clark, W., Leach, J.D., Jackstadt, R., Nixon, C., Huels, D.J., et al. (2019). Loss of BCL9/9l suppresses Wnt driven tumourigenesis in models that recapitulate human cancer. Nat Commun 10, 723. 10.1038/s41467-019-08586-3.

20. De Lau, W., Peng, W.C., Gros, P., and Clevers, H. (2014). The R-spondin/Lgr5/Rnf43 module: regulator of Wnt signal strength. Genes Dev. 28, 305–316. 10.1101/gad.235473.113.

21. Nusse, R., and Clevers, H. (2017). Wnt/β-Catenin Signaling, Disease, and Emerging Therapeutic Modalities. Cell 169, 985–999. 10.1016/j.cell.2017.05.016.

22. Molenaar, M., Van De Wetering, M., Oosterwegel, M., Peterson-Maduro, J., Godsave, S., Korinek, V., Roose, J., Destrée, O., and Clevers, H. (1996). XTcf-3 Transcription Factor Mediates β-Catenin-Induced Axis Formation in Xenopus Embryos. Cell 86, 391–399. 10.1016/S0092-8674(00)80112-9.

23. Cadigan, K.M., and Waterman, M.L. (2012). TCF/LEFs and Wnt Signaling in the Nucleus. Cold Spring Harbor Perspectives in Biology 4, a007906–a007906. 10.1101/cshperspect.a007906.

24. Stamos, J.L., and Weis, W.I. (2013). The -Catenin Destruction Complex. Cold Spring Harbor Perspectives in Biology 5, a007898–a007898. 10.1101/cshperspect.a007898.

25. Rubinfeld, B., Albert, I., Porfiri, E., Fiol, C., Munemitsu, S., and Polakis, P. (1996). Binding of GSK3β to the APC-β-Catenin Complex and Regulation of Complex Assembly. Science 272, 1023–1026. 10.1126/science.272.5264.1023.

26. Wagner, F.F., Benajiba, L., Campbell, A.J., Weïwer, M., Sacher, J.R., Gale, J.P., Ross, L., Puissant, A., Alexe, G., Conway, A., et al. (2018). Exploiting an Asp-Glu “switch” in glycogen synthase kinase 3 to design paralog-selective inhibitors for use in acute myeloid leukemia. Sci Transl Med 10, eaam8460. 10.1126/scitranslmed.aam8460.

27. Cohen, P., and Goedert, M. (2004). GSK3 inhibitors: development and therapeutic potential. Nat Rev Drug Discov 3, 479–487. 10.1038/nrd1415.

28. Vidri, R.J., and Fitzgerald, T.L. (2020). GSK-3: An important kinase in colon and pancreatic cancers. Biochimica et Biophysica Acta (BBA) - Molecular Cell Research 1867, 118626. 10.1016/j.bbamcr.2019.118626.

29. Engler, T.A., Henry, J.R., Malhotra, S., Cunningham, B., Furness, K., Brozinick, J., Burkholder, T.P., Clay, M.P., Clayton, J., Diefenbacher, C., et al. (2004). Substituted 3-Imidazo[1,2-a]pyridin-3-yl-4-(1,2,3,4-tetrahydro-[1,4]diazepino-[6,7,1-hi]indol-7-yl)pyrrole-2,5-diones as Highly Selective and Potent Inhibitors of Glycogen Synthase Kinase-3. J. Med. Chem. 47, 3934–3937. 10.1021/jm049768a.

30. Staal, F.J.T., Noort, M. van, Strous, G.J., and Clevers, H.C. (2002). Wnt signals are transmitted through N-terminally dephosphorylated β-catenin. EMBO Rep 3, 63–68. 10.1093/embo-reports/kvf002.

31. Miyoshi, H., and Stappenbeck, T.S. (2013). In vitro expansion and genetic modification of gastrointestinal stem cells in spheroid culture. Nat Protoc 8, 2471–2482. 10.1038/nprot.2013.153.

32. VanDussen, K.L., Sonnek, N.M., and Stappenbeck, T.S. (2019). L-WRN conditioned medium for gastrointestinal epithelial stem cell culture shows replicable batch-to-batch activity levels across multiple research teams. Stem Cell Research 37, 101430. 10.1016/j.scr.2019.101430.

33. Cohen, P., and Goedert, M. (2004). GSK3 inhibitors: development and therapeutic potential. Nat Rev Drug Discov 3, 479–487. 10.1038/nrd1415.

34. An, W.F., Germain, A.R., Bishop, J.A., Nag, P.P., Metkar, S., Ketterman, J., Walk, M., Weiwer, M., Liu, X., Patnaik, D., et al. (2010). Discovery of Potent and Highly Selective Inhibitors of GSK3b. In Probe Reports from the NIH Molecular Libraries Program (National Center for Biotechnology Information (US)).

35. Zamek-Gliszczynski, M.J., Abraham, T.L., Alberts, J.J., Kulanthaivel, P., Jackson, K.A., Chow, K.H., McCann, D.J., Hu, H., Anderson, S., Furr, N.A., et al. (2013). Pharmacokinetics, Metabolism, and Excretion of the Glycogen Synthase Kinase-3 Inhibitor LY2090314 in Rats, Dogs, and Humans: A Case Study in Rapid Clearance by Extensive Metabolism with Low Circulating Metabolite Exposure. Drug Metab Dispos 41, 714–726. 10.1124/dmd.112.048488.

36. Enos, M.D., Gavagan, M., Jameson, N., Zalatan, J.G., and Weis, W.I. (2024). Structural and functional effects of phosphopriming and scaffolding in the kinase GSK-3β. Science Signaling 17, eado0881. 10.1126/scisignal.ado0881.

37. Ye, S., Tan, L., Yang, R., Fang, B., Qu, S., Schulze, E.N., Song, H., Ying, Q., and Li, P. (2012). Pleiotropy of Glycogen Synthase Kinase-3 Inhibition by CHIR99021 Promotes Self-Renewal of Embryonic Stem Cells from Refractory Mouse Strains. PLOS ONE 7, e35892. 10.1371/journal.pone.0035892.

38. Kim, H., Wu, J., Ye, S., Tai, C.-I., Zhou, X., Yan, H., Li, P., Pera, M., and Ying, Q.-L. (2013). Modulation of β-catenin function maintains mouse epiblast stem cell and human embryonic stem cell self-renewal. Nat Commun 4, 2403. 10.1038/ncomms3403.

39. Ying, Q.-L., Wray, J., Nichols, J., Batlle-Morera, L., Doble, B., Woodgett, J., Cohen, P., and Smith, A. (2008). The ground state of embryonic stem cell self-renewal. Nature 453, 519–523. 10.1038/nature06968.

40. Yin, X., Farin, H.F., van Es, J.H., Clevers, H., Langer, R., and Karp, J.M. (2014). Niche-independent high-purity cultures of Lgr5+ intestinal stem cells and their progeny. Nat Methods 11, 106–112. 10.1038/nmeth.2737.

41. Chen, H., and Langer, R. (1998). Oral particulate delivery: status and future trends. Advanced Drug Delivery Reviews 34, 339–350. 10.1016/S0169-409X(98)00047-7.

42. Enteric Infections in Young Children are Associated with Environmental Enteropathy and Impaired Growth - George - 2018 - Tropical Medicine & International Health - Wiley Online Library https://onlinelibrary.wiley.com/doi/full/10.1111/tmi.13002.

43. Teodósio, N.R., Lago, E.S., Romani, S.A., and Guedes, R.C. (1990). A regional basic diet from northeast Brazil as a dietary model of experimental malnutrition. Arch Latinoam Nutr 40, 533–547.

44. Korpe, P.S., and Petri, W.A. (2012). Environmental enteropathy: critical implications of a poorly understood condition. Trends in Molecular Medicine 18, 328–336. 10.1016/j.molmed.2012.04.007.

45. Keusch, G.T., Denno, D.M., Black, R.E., Duggan, C., Guerrant, R.L., Lavery, J.V., Nataro, J.P., Rosenberg, I.H., Ryan, E.T., Tarr, P.I., et al. (2014). Environmental Enteric Dysfunction: Pathogenesis, Diagnosis, and Clinical Consequences. Clinical Infectious Diseases 59, S207–S212. 10.1093/cid/ciu485.

46. Ueno, P.M., Oriá, R.B., Maier, E.A., Guedes, M., de Azevedo, O.G., Wu, D., Willson, T., Hogan, S.P., Lima, A.A.M., Guerrant, R.L., et al. (2011). Alanyl-glutamine promotes intestinal epithelial cell homeostasis in vitro and in a murine model of weanling undernutrition. Am J Physiol Gastrointest Liver Physiol 301, G612–G622. 10.1152/ajpgi.00531.2010.

47. Bonnet, C., Brahmbhatt, A., Deng, S.X., and Zheng, J.J. Wnt signaling activation: targets and therapeutic opportunities for stem cell therapy and regenerative medicine. RSC Chem Biol 2, 1144– 1157. 10.1039/d1cb00063b.

48. Goto, N., Goto, S., Imada, S., Hosseini, S., Deshpande, V., and Yilmaz, Ö.H. (2022). Lymphatics and fibroblasts support intestinal stem cells in homeostasis and injury. Cell Stem Cell 29, 1246–1261.e6. 10.1016/j.stem.2022.06.013.

49. Willert, K., Brown, J.D., Danenberg, E., Duncan, A.W., Weissman, I.L., Reya, T., Yates, J.R., and Nusse, R. (2003). Wnt proteins are lipid-modified and can act as stem cell growth factors. Nature 423, 448–452. 10.1038/nature01611.

50. Tüysüz, N., van Bloois, L., van den Brink, S., Begthel, H., Verstegen, M.M.A., Cruz, L.J., Hui, L., van der Laan, L.J.W., de Jonge, J., Vries, R., et al. (2017). Lipid-mediated Wnt protein stabilization enables serum-free culture of human organ stem cells. Nat Commun 8, 14578. 10.1038/ncomms14578.

51. Veeman, M.T., Slusarski, D.C., Kaykas, A., Louie, S.H., and Moon, R.T. (2003). Zebrafish prickle, a modulator of noncanonical Wnt/Fz signaling, regulates gastrulation movements. Curr Biol 13, 680– 685. 10.1016/s0960-9822(03)00240-9.

52. Jayne, D.G., Fook, S., Loi, C., and Seow-Choen, F. (2002). Peritoneal carcinomatosis from colorectal cancer. BJS (British Journal of Surgery) 89, 1545–1550. 10.1046/j.1365-2168.2002.02274.x.

53. Colon Cancer, Version 2.2021, NCCN Clinical Practice Guidelines in Oncology in: Journal of the National Comprehensive Cancer Network Volume 19 Issue 3 (2021) https://jnccn.org/view/journals/jnccn/19/3/article-p329.xml?ArticleBodyColorStyles=full%20html.

54. Riihimäki, M., Hemminki, A., Sundquist, J., and Hemminki, K. (2016). Patterns of metastasis in colon and rectal cancer. Sci Rep 6, 29765. 10.1038/srep29765.

55. Moser, A.R., Pitot, H.C., and Dove, W.F. (1990). A Dominant Mutation That Predisposes to Multiple Intestinal Neoplasia in the Mouse. Science 247, 322–324. 10.1126/science.2296722.

56. Love, M.I., Huber, W., and Anders, S. (2014). Moderated estimation of fold change and dispersion for RNA-seq data with DESeq2. Genome Biol 15, 550. 10.1186/s13059-014-0550-8.

57. Yang, Y., and Mlodzik, M. (2015). Wnt-Frizzled/Planar Cell Polarity Signaling: Cellular Orientation by Facing the Wind (Wnt). Annual Review of Cell and Developmental Biology 31, 623–646. 10.1146/annurev-cellbio-100814-125315.

58. Wan, C., Mahara, S., Sun, C., Doan, A., Chua, H.K., Xu, D., Bian, J., Li, Y., Zhu, D., Sooraj, D., et al. (2021). Genome-scale CRISPR-Cas9 screen of Wnt/β-catenin signaling identifies therapeutic targets for colorectal cancer. Sci Adv 7, eabf2567. 10.1126/sciadv.abf2567.

59. Shi, J., and Wei, L. (2007). Rho kinase in the regulation of cell death and survival. Arch. Immunol. Ther. Exp. 55, 61–75. 10.1007/s00005-007-0009-7.

60. Croft, D.R., Coleman, M.L., Li, S., Robertson, D., Sullivan, T., Stewart, C.L., and Olson, M.F. (2005). Actin-myosin–based contraction is responsible for apoptotic nuclear disintegration. J Cell Biol 168, 245–255. 10.1083/jcb.200409049.

61. Marino, S., Vooijs, M., Gulden, H. van der, Jonkers, J., and Berns, A. (2000). Induction of medulloblastomas in p53-null mutant mice by somatic inactivation of Rb in the external granular layer cells of the cerebellum. Genes Dev. 14, 994–1004. 10.1101/gad.14.8.994.

62. Jackson, E.L., Willis, N., Mercer, K., Bronson, R.T., Crowley, D., Montoya, R., Jacks, T., and Tuveson, D.A. (2001). Analysis of lung tumor initiation and progression using conditional expression of oncogenic K-ras. Genes Dev. 15, 3243–3248. 10.1101/gad.943001.

63. Madisen, L., Zwingman, T.A., Sunkin, S.M., Oh, S.W., Zariwala, H.A., Gu, H., Ng, L.L., Palmiter, R.D., Hawrylycz, M.J., Jones, A.R., et al. (2010). A robust and high-throughput Cre reporting and characterization system for the whole mouse brain. Nat Neurosci 13, 133–140. 10.1038/nn.2467.

64. Kuraguchi, M., Wang, X.-P., Bronson, R.T., Rothenberg, R., Ohene-Baah, N.Y., Lund, J.J., Kucherlapati, M., Maas, R.L., and Kucherlapati, R. (2006). Adenomatous Polyposis Coli (APC) Is Required for Normal Development of Skin and Thymus. PLOS Genetics 2, e146. 10.1371/journal.pgen.0020146.

65. El Marjou, F., Janssen, K.-P., Hung-Junn Chang, B., Li, M., Hindie, V., Chan, L., Louvard, D., Chambon, P., Metzger, D., and Robine, S. (2004). Tissue-specific and inducible Cre-mediated recombination in the gut epithelium. genesis 39, 186–193. 10.1002/gene.20042.

66. Sánchez-Rivera, F.J., Papagiannakopoulos, T., Romero, R., Tammela, T., Bauer, M.R., Bhutkar, A., Joshi, N.S., Subbaraj, L., Bronson, R.T., Xue, W., et al. (2014). Rapid modelling of cooperating genetic events in cancer through somatic genome editing. Nature 516, 428–431. 10.1038/nature13906.

67. Dow, L.E., O’Rourke, K.P., Simon, J., Tschaharganeh, D.F., van Es, J.H., Clevers, H., and Lowe, S.W. (2015). Apc Restoration Promotes Cellular Differentiation and Reestablishes Crypt Homeostasis in Colorectal Cancer. Cell 161, 1539–1552. 10.1016/j.cell.2015.05.033.

68. Cheng, C.-W., Biton, M., Haber, A.L., Gunduz, N., Eng, G., Gaynor, L.T., Tripathi, S., Calibasi-Kocal, G., Rickelt, S., Butty, V.L., et al. (2019). Ketone Body Signaling Mediates Intestinal Stem Cell Homeostasis and Adaptation to Diet. Cell 178, 1115–1131.e15. 10.1016/j.cell.2019.07.048.

69. Patro, R., Duggal, G., Love, M.I., Irizarry, R.A., and Kingsford, C. (2017). Salmon provides fast and bias-aware quantification of transcript expression. Nat Methods 14, 417–419. 10.1038/nmeth.4197.

70. Soneson, C., Love, M.I., and Robinson, M.D. (2016). Differential analyses for RNA-seq: transcript-level estimates improve gene-level inferences. F1000Res 4, 1521. 10.12688/f1000research.7563.2.

71. Ge, S.X., Son, E.W., and Yao, R. (2018). iDEP: an integrated web application for differential expression and pathway analysis of RNA-Seq data. BMC Bioinformatics 19, 534. 10.1186/s12859-018-2486-6.

72. Langmead, B., and Salzberg, S.L. (2012). Fast gapped-read alignment with Bowtie 2. Nat Methods 9, 357–359. 10.1038/nmeth.1923.

73. Li, H., Handsaker, B., Wysoker, A., Fennell, T., Ruan, J., Homer, N., Marth, G., Abecasis, G., Durbin, R., and 1000 Genome Project Data Processing Subgroup (2009). The Sequence Alignment/Map format and SAMtools. Bioinformatics 25, 2078–2079. 10.1093/bioinformatics/btp352.

74. Quinlan, A.R., and Hall, I.M. (2010). BEDTools: a flexible suite of utilities for comparing genomic features. Bioinformatics 26, 841–842. 10.1093/bioinformatics/btq033.

75. Zhang, Y., Liu, T., Meyer, C.A., Eeckhoute, J., Johnson, D.S., Bernstein, B.E., Nusbaum, C., Myers, R.M., Brown, M., Li, W., et al. (2008). Model-based analysis of ChIP-Seq (MACS). Genome Biol 9, R137. 10.1186/gb-2008-9-9-r137.

